# Cognition emerges from phase dynamics of intrinsic coordination

**DOI:** 10.64898/2026.03.26.714488

**Authors:** Youngjo Song, Jiayu Chen, Vince D. Calhoun, Armin Iraji

## Abstract

The brain generates diverse cognitive states while maintaining a stable functional architecture, a duality that remains difficult to reconcile. Prevailing views assume that flexible cognition necessitates correspondingly flexible architecture with different tasks demanding distinct reconfigurations, leaving the coexistence of stability and flexibility unexplained. Here we introduce the intrinsic network flow (INF) framework as a complementary view. This framework is built on temporally coordinated signal flows across brain networks that constitute a universal scaffold, stable across diverse cognitive states and common across individuals. We show that a wide range of task-evoked activation patterns can be reconstructed by modulating only the temporal phase alignment of these flows, whose fixed structure determines functional connectivity topology, gradients, and large-scale networks, thereby preserving these properties across task states and reconciling flexibility with stability. This situates resting-state and task-state dynamics within a unified framework and suggests a generative relationship from flow-like dynamics to the full landscape of resting-state and task-state phenomena. Crucially, phase information achieved 89.9% accuracy in classifying 23 cognitive tasks, outperforming amplitude- or activation-based markers. These findings reframe task-evoked activation and deactivation as interference among concurrent flows. Together, our work indicates that a key variable for cognition is when intrinsic dynamics align in time, not where or how much the brain activates.

## Article

How biological systems achieve flexible function under largely stable architecture remains a fundamental question [1, 2]. In particular, brain dynamics exhibit a pronounced stability– flexibility duality: intrinsic functional connectivity (FC) topology is highly conserved across tasks [3-5], reflecting stable patterns of large-scale communication and integration, whereas task-evoked activation exhibits substantial spatial variability [4], consistent with context-dependent engagement of neural populations. Despite extensive empirical characterization, how such context-dependent computations arise without disrupting the underlying organizational scaffold remains unresolved, maintaining a dichotomy between resting-state and task-based approaches in neuroscience.

This duality presents a specific paradox: regions that appear unrelated or even antagonistic in terms of intrinsic FC—being segregated or anti-correlated at rest (e.g., default mode versus executive networks [6])—are nevertheless frequently co-recruited during complex tasks (e.g., [7, 8]). Prevailing accounts resolve this through task-induced departures from intrinsic organization: e.g., dynamic network reconfiguration [9-11] or transient brain states that are obscured by time-averaged FC estimates [12-15], which are potentially driven by flexible inputs [16, 17]. However, these frameworks lack a principled mechanism for preserving FC topology across tasks because they rely on departures from the intrinsic scaffold. Moreover, they face a scaling problem: accommodating diverse task demands leads to a combinatorial explosion in the space of required configurations, inputs, or states.

Notably, it has been suggested that task-evoked activation is not a simple additive response to resting-state organization, but rather a structured or limited expression of underlying intrinsic dynamics [5, 18]. Consistent with this view, resting-state and task-state trajectories can be embedded within a shared low-dimensional manifold [19], and task-evoked activity patterns align closely with topographic maps of statistically independent resting-state activity [20].

Although these observations point toward a unified principle linking rest and task dynamics, they do not specify the generative mechanism by which the stable intrinsic architecture produces diverse task-dependent patterns. Here, we argue that resolving this gap, together with the aforementioned antagonist co-recruitment paradox, requires moving beyond extensions of existing models to a new and complementary conceptual level for describing large-scale brain dynamics, which we formalize as the intrinsic network flow (INF) framework.

### Intrinsic Network Flow (INF)

We posit that large-scale brain dynamics are governed by a stable spatiotemporal coordination structure—a set of signal flows across networks, which we term INFs—that is conserved across resting and task states. Diverse cognitive states emerge not from reconfiguring these flows or simply modulating regional activation amplitudes, but through the *phase dynamics* of the flows. Specifically, modulation of the relative temporal alignment between concurrent flows determines which networks are engaged at any given moment, without altering the structure of flows themselves.

This framework is motivated by converging observations that resting-state brain activity exhibits reproducible spatiotemporal patterns, including quasi-periodic patterns [21, 22] and wave-like propagation [23-26]. These patterns have been shown to potentially serve as generative sources of diverse functional organizational features, including average/time-varying FC topology, functional hierarchy captured by gradient mapping, and network modules [27, 28], and their morphologies are largely preserved during task performance [21, 28]. Accordingly, the present framework treats spatial and temporal organization as fundamentally coupled rather than separable [29]. This contrasts with existing dynamical frameworks, which embed spatial activation patterns into a low-dimensional space and analyze temporal evolution within that space (e.g., [17, 30, 31]), thereby treating the two dimensions as at least partially independent.

#### Formulation of the INF framework

We define “flow” as a temporally ordered coordination pattern across spatially distributed intrinsic networks (INs), specifying which networks participate and in what sequence. Each flow is formalized as an INF mode: a pair of complex-conjugate vectors 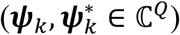, where *Q* is the number of INs (Figure 1a; illustrated with a three-network toy example). In this representation, the magnitude of each element (|(***ψ***_*k*_)_*q*_|, where *q* indexes IN) quantifies the *degree of network participation*, while its argument (arg[(***ψ***_*k*_)_*q*_]) encodes the network’s relative position within the *coordinated flow sequence*—the fixed temporal ordering of leads and lags that defines the flow’s identity. Thus, ***ψ***_*k*_ concisely describes “*who participates*” and “*in what order and by how much delay*.” Crucially, these modes are identified at the group level and are conserved across individuals and cognitive states, forming a universal scaffold for brain dynamics.

**Figure 1.**
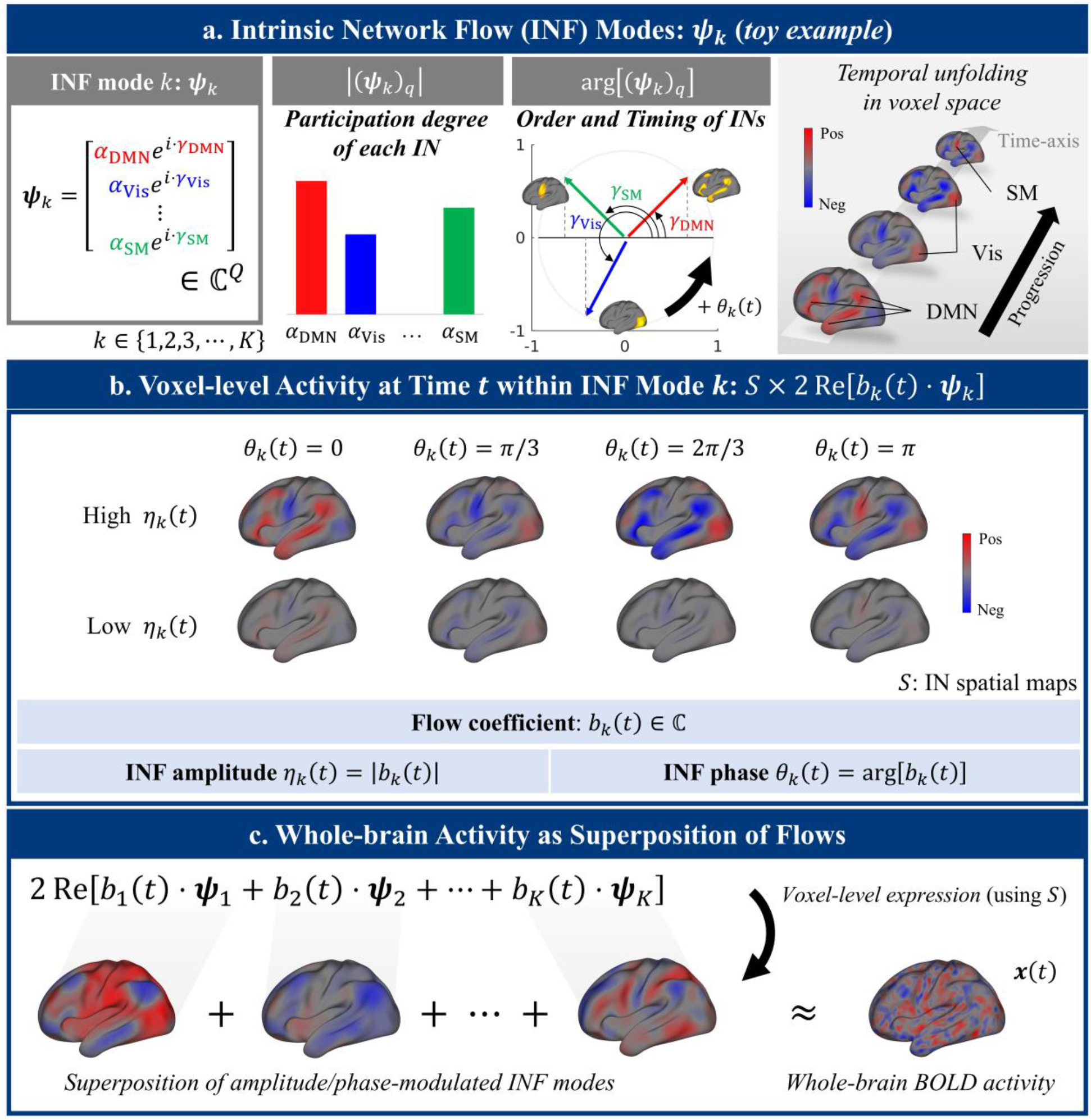
Formulation of the INF framework. **(a)** An intrinsic network flow (INF) mode ***ψ***_*k*_ is a complex-valued vector over *Q* intrinsic networks (INs). The magnitude of each element (|(***ψ***_*k*_)_*q*_|) encodes the participation degree of the corresponding IN (shown on the bar graph, where *q* indexes IN). The argument of each element (arg[(***ψ***_*k*_)_*q*_]) encodes flow sequence (shown on the unit circle): the fixed temporal ordering of phase leads and lags between networks. A toy example with three participating INs (DMN, visual, and SM) illustrates how these components unfold as a spatiotemporal flow in voxel space (right). As the INF phase *θ*_*k*_(*t*) increases, the complex representation of each network rotates counter-clockwise in the complex plane (e.g., *γ*_DMN_ + *θ*_*k*_(*t*) → *γ*_DMN_ + *θ*_*k*_(*t* + 1) ; see arg[(***ψ***_*k*_)_*q*_] diagram). The observed activity is the real-valued projection onto the real axis. Thus, in this toy example, the mode flows in a sequence of DMN, Vis, then SM. **(b)** At each time point *t*, the network-level activity of mode *k* is given by 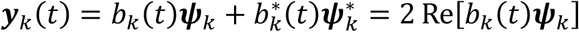, where *b*_*k*_(*t*) is the complex-valued flow coefficient. Cortical patterns shown here are obtained by projecting network-level activity through IN spatial maps (*S* × ***y***_*k*_(*t*)). The INF amplitude (*η*_*k*_ = |*b*_*k*_(*t*)|) determines the overall strength of engagement of this flow, while the INF phase (*θ*_*k*_(*t*) = arg[*b*_*k*_(*t*)]) determines the current position within the flow’s temporal cycle, specifying which networks are engaged at that instant. Rows show the effect of amplitude (high vs. low); columns show the effect of phase advancing from 0 to *π*. Because shifting the phase changes which networks are engaged at a given instant, a single mode can produce diverse spatial activation patterns. **(c)** Whole-brain BOLD activity ***x***(*t*) is modeled as the superposition of all INF modes projected into voxel space (***x***(*t*) = *S* ⋅ ∑_*k*_ ***y***_*k*_(*t*) = *S* ⋅ ∑_*k*_ 2 Re[*b*_*k*_(*t*)***ψ***_*k*_]), where the modes themselves (***ψ***_*k*_) remain fixed and only the flow coefficients *b*_*k*_(*t*) vary. DMN: default mode network. Vis: visual network. SM: somatomotor network.

#### Flow coefficients and instantaneous brain activity

The moment-to-moment expression of each INF mode is governed by a complex-valued *flow coefficient b*_*k*_(*t*), which determines the mode’s network-level activity at time t as 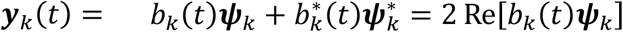 (Figure 1b). Real-valued reconstruction is guaranteed by conjugate pairing of the flow coefficients and mode vectors. Multiple modes coexist, and their superposition produces the spatiotemporal complexity of whole-brain dynamics at the network level (***y***(*t*) = ∑_*k*_ ***y***_*k*_(*t*) = ∑_*k*_ 2 Re[*b*_*k*_(*t*)***ψ***_*k*_]), which maps to whole-brain BOLD activity through IN spatial maps *S* (***x***(*t*) = *S* ⋅ ***y***(*t*); Figure 1c).

Within this formulation, *INF amplitude* (*η*_*k*_(*t*) = |*b*_*k*_(*t*)|) quantifies how strongly the flow is expressed at time t, while *INF phase* (*θ*_*k*_(*t*) = arg[*b*_*k*_(*t*)]) specifies the flow’s current position within its temporal cycle. It is critical to distinguish between two types of timing in this framework. (1) The flow sequence (arg[(***ψ***_*k*_)_*q*_]) is the fixed ordering of network engagements that defines a given mode’s identity. (2) The INF phase (*θ*_*k*_(*t*)), by contrast, is a dynamic variable that quantifies the timing of each mode as it evolves over time. Our results suggest that the *INF phase* (*θ*_*k*_(*t*)) is a crucial parameter for characterizing cognitive states: shifting the phases of concurrent INF modes changes which networks are engaged at a given instant, such that diverse states are realized without requiring any alteration of the underlying coordination structures (i.e., INF modes ***ψ***_*k*_).

#### Spontaneous evolution of flows

Even in the absence of task or external demands, the brain is never static but rather continuously evolves. To capture this, each flow coefficient evolves through multiplication by a characteristic complex scalar *d*_*k*_—the intrinsic evolution rate—advancing the INF phase at a rate intrinsic to the mode (i.e., *b*_*k*_(*t* + 1) = *d*_*k*_ ⋅ *b*_*k*_(*t*)) and producing the spontaneous temporal progression observed at rest (Supplementary Figure 1). This spontaneous evolution does not cease during task performance—task-related phase modulation acts on top of the ongoing intrinsic progression, rather than replacing it. This property provides the basis for estimating INF modes from fMRI data; the procedure is summarized in Figure 2 with full details in Methods.

**Figure 2.**
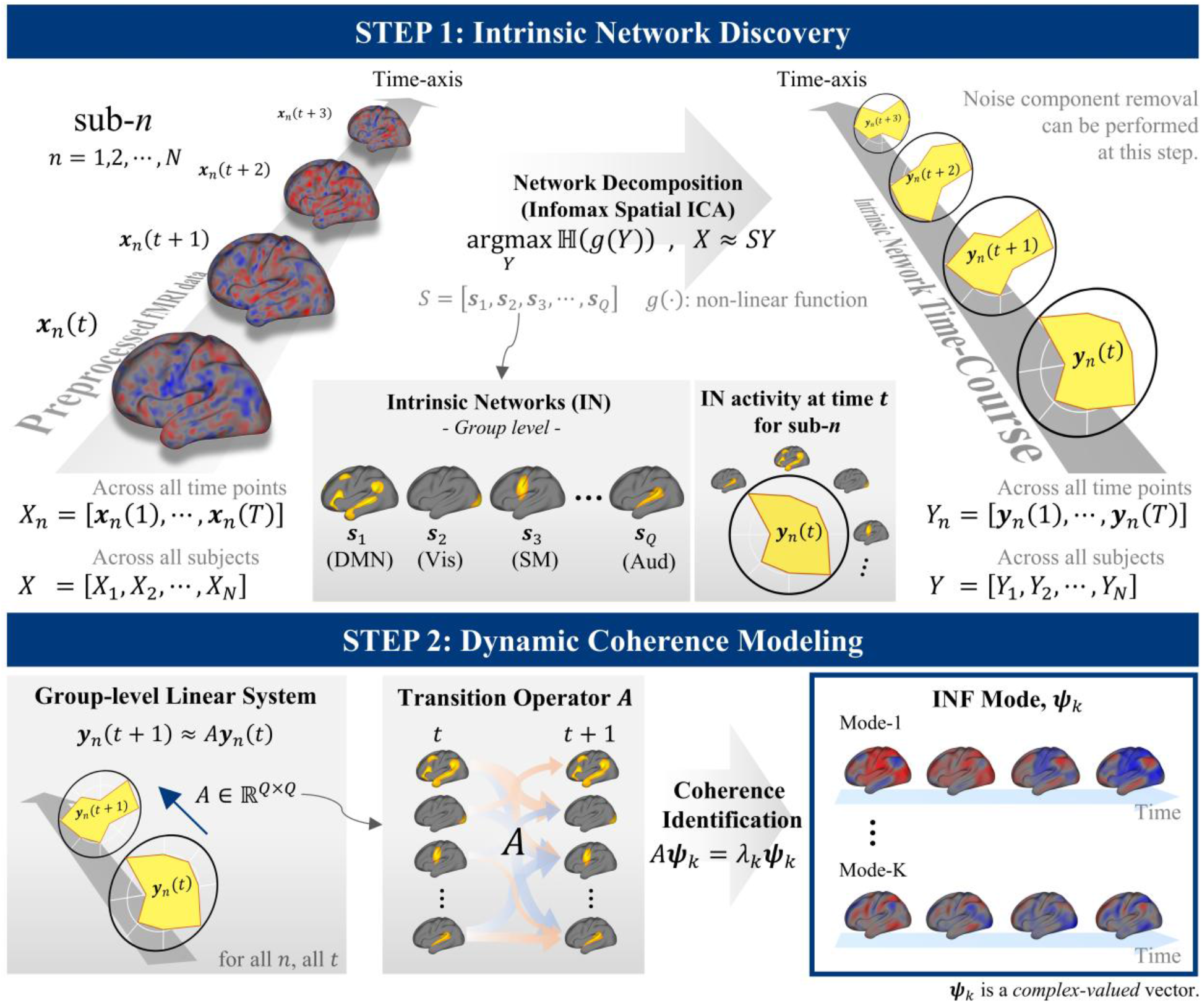
INF mode estimation procedure. **(STEP 1)** Intrinsic network (IN) discovery: group ICA decomposes all subjects’ BOLD signals into *Q* group-level IN spatial maps (*S*) and their associated time courses (*Y*). **(STEP 2)** Dynamic coherence modeling: the temporal evolution of IN time courses is modeled as a group-level linear system ***y***_*n*_(*t* + 1) = *A****y***_*n*_(*t*), estimated across all subjects and time points. Eigendecomposition of the transition operator *A* yields INF modes ***ψ***_*k*_ —complex-valued eigenvectors representing coherent flow structures. Individual variability is subsequently characterized through spatial and temporal fingerprinting (Supplementary Figure 2). See Methods for implementation details.

## Results

### Predictive Validation of the Coordination Structure

Central to the INF framework is a stable spatiotemporal coordination structure underlying brain dynamics (i.e., INF modes). We therefore tested whether such a structure can be reliably identified and how many modes are required to capture it. To this end, INF modes were extracted from a training sample (*N* = 105, human connectome project (HCP) dataset; see Methods for detailed procedure) and evaluated on their ability to predict future BOLD signals in an independent test sample (*N* = 105), benchmarked against a first-order autoregressive (AR(1)) null model and subject-specific coordination structures (i.e., flows) estimated from each individual’s data alone.

Across all tested dimensionalities (*Q* = 10–100), prediction performance (*R*^*2*^) significantly outperformed the null model (p < 1.0 × 10^-72^; Figure 3), indicating that the extracted flows capture spatiotemporal organization not reducible to simple temporal autocorrelation. Notably, across all tested dimensionalities, INF modes estimated at the group level yielded higher predictive accuracy for individual BOLD signals than modes estimated from subject-specific data (Figure 3). Although this improvement may reflect the larger amount of data underlying the group-level estimates, the result nonetheless suggests that the brain’s flow-like coordination structure is a common, stable scaffold that is conserved across individuals, since otherwise group-level modes would harm the accuracy.

**Figure 3.**
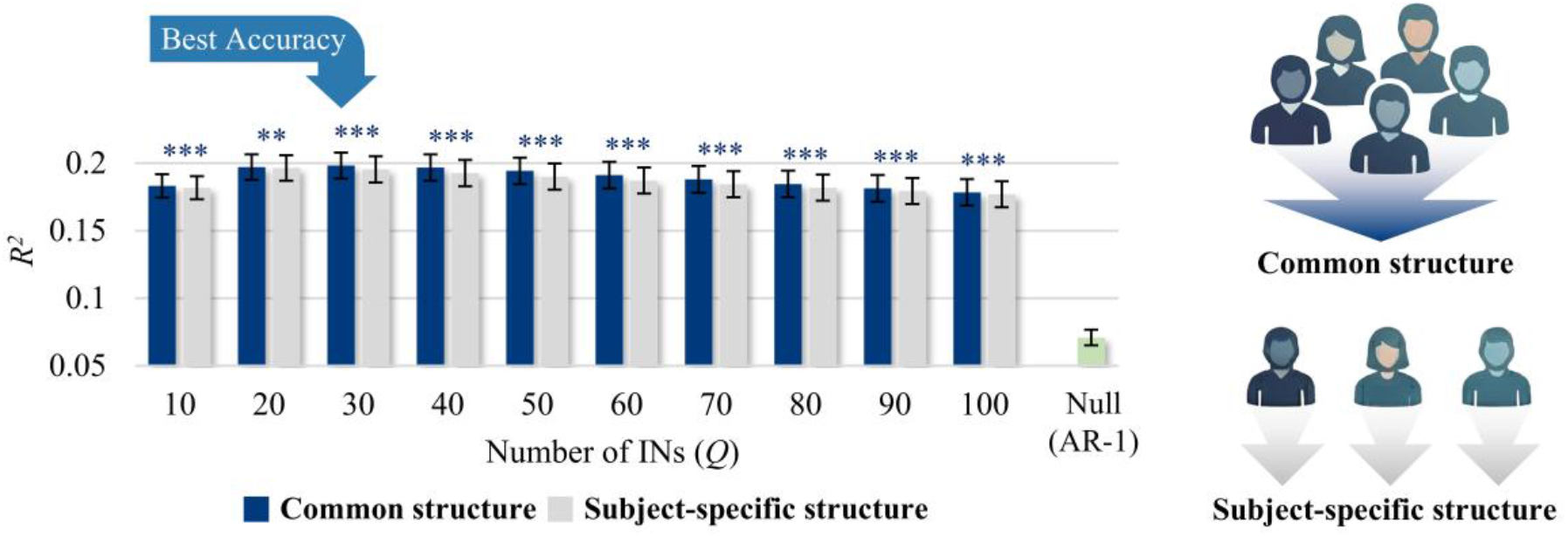
Predictive validation of the coordination structure (i.e., INF modes). Cross-validated prediction performance (*R*^*2*^) of future BOLD signals using a group-level (common) coordination structure (i.e., group-level INF modes; dark blue) versus subject-specific structures estimated from each individual’s data alone (i.e., subject-specific INF mode; grey), across IN dimensionalities (Q = 10–100). The AR(1) null model (green) is shown for comparison. Paired t-tests compared common and subject-specific structures (*N* = 105). Asterisks indicate significance (**p < 0.01; ***p < 0.001); dark blue asterisks indicate the common structure outperformed the subject-specific structure, and grey asterisks indicate the opposite. Error bars denote 95% confidence intervals.

Prediction accuracy peaked at a dimensionality of 27 networks with group-level modes (corresponding to 13 INF mode pairs; see Methods; Supplementary Figure 3a), a scale that was adopted for all subsequent analyses. All findings were replicated in an independent sample in HCP dataset (*N* = 210; see Supplementary Result 1).

### INF 1: The principal flow recapitulates dominant whole-brain functional organization

We next asked what these flows encode. To maximize robustness for characterizing flow structure, we estimated INF modes from the full HCP REST1 dataset (*N* = 951) using the dimensionality identified above (*Q* = 27). The modes identified here were largely reproduced in the REST2 retest session (*N* = 884; Supplementary Result 2). If INF modes capture the brain’s fundamental spatiotemporal scaffold, their structure should be systematically related to known macroscale functional organization.

INF mode 1—the mode with the largest overall amplitude—reflects a coordinated temporal progression between sensorimotor systems (unimodal areas; visual, somatomotor, and auditory areas) and association cortex (transmodal areas), predominantly the default mode network (DMN) (Figure 4a; Supplementary Video 1). Its flow sequence is organized primarily along the sensorimotor–association (SA) hierarchy, with somatomotor regions slightly leading visual regions, introducing an additional timing offset orthogonal to this primary progression (Figure 4b, left). Notably, the voxel-level projection of flow sequence encapsulate the dominant axes of cortical FC organization—the SA hierarchy (FC gradient 1) and the visual–somatomotor (VS) axis (FC gradient 2) [32] (Figure 4b, right). Quantitatively, the orthogonal components of the complex flow sequence map align remarkably well with these gradients. The cosine component correlated strongly with the SA axis (R = 0.88, p < 0.001, spin test; Figure 4c) and the sine component with the VS axis (R = 0.64, p = 0.008, spin test; Figure 4d). This indicates that the two principal cortical gradients are not independent organizational axes, but rather orthogonal projections of a single temporally coordinated flow.

**Figure 4.**
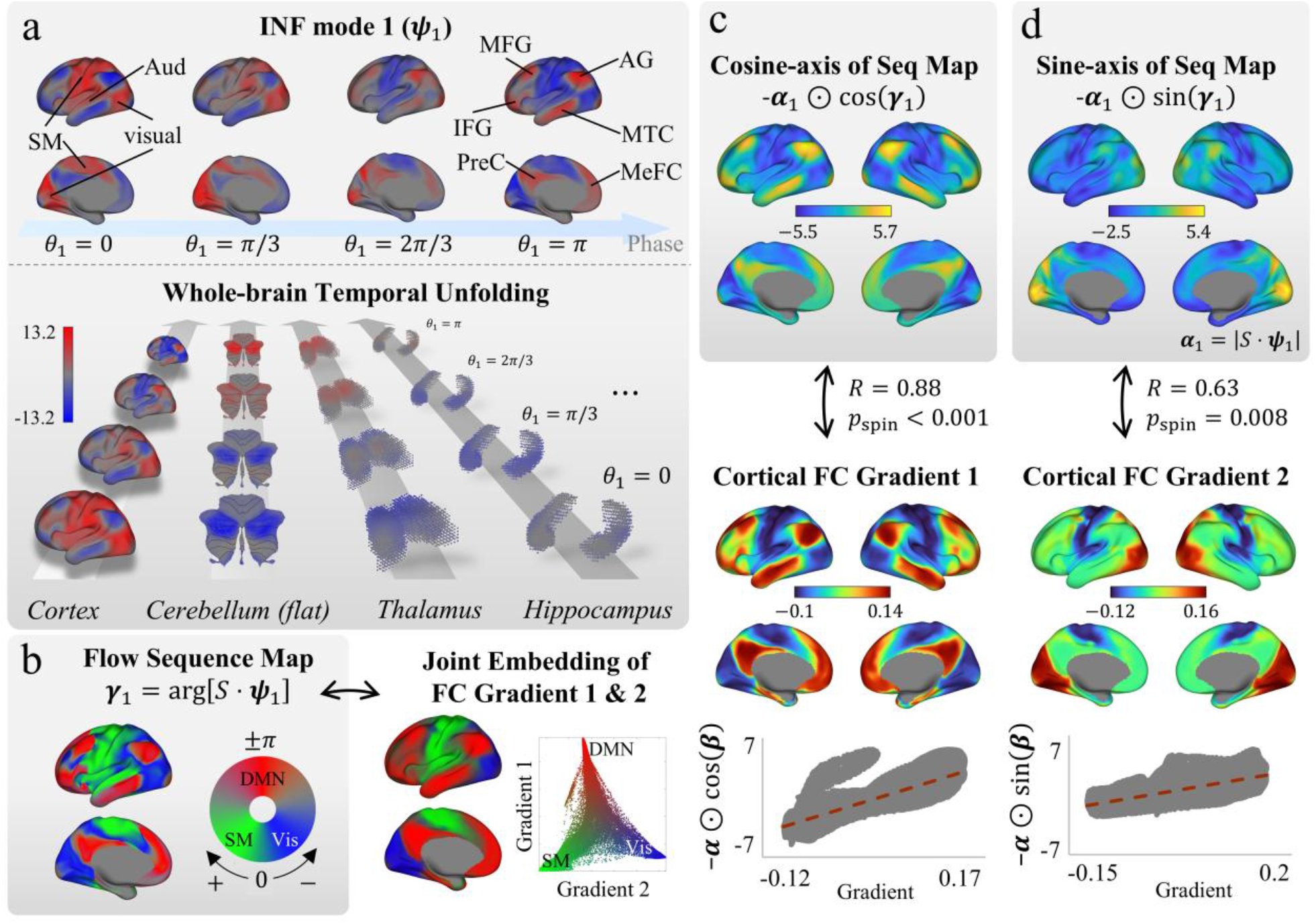
The principal flow recapitulates dominant cortical functional organization. **(a)** INF mode 1 (***ψ***_1_), the mode with the largest overall amplitude, describes a coordinated temporal progression between sensorimotor systems and association cortex. Top: cortical surface activity at four INF phase values (*θ*_1_ = 0, π/3, 2π/3, π), showing the sequential engagement of sensorimotor regions followed by association areas. Bottom: whole-brain temporal unfolding across cortex, cerebellum (flatmap), thalamus, and hippocampus. Voxel-level patterns were obtained by projecting the INF mode vector through the group-level IN spatial maps *S* (i.e., *S* · ***ψ***_1_). **(b)** Left (grey box): the flow sequence map (***γ***_1_ = arg[*S* · ***ψ***_1_]) displayed on the cortical surface, indicating that the flow progresses primarily along the sensorimotor–association (SA) hierarchy, with an additional timing offset between visual and somatomotor systems. Right: joint embedding of cortical FC gradients 1 and 2, showing a corresponding spatial arrangement of DMN, visual, and SM regions. The scatter plot displays gradient 1 versus gradient 2 values for individual vertices. **(c)** Top (grey box): The cosine-axis projection of the flow sequence map (−***α***_1_ ⊙ cos(***γ***_1_), modulated by magnitude ***α***_1_ = |*S* · ***ψ***_1_|; ⊙ denotes element-wise multiplication) compared with cortical FC gradient 1 (SA hierarchy). Middle: cortical FC gradient 1. Bottom: vertex-wise scatter plot (R = 0.88, p < 0.001, spin test). **(d)** Top (grey box): The sine-axis projection (−***α***_1_ ⊙ sin(***γ***_1_)) compared with cortical FC gradient 2 (visual–somatomotor axis). Middle: cortical FC gradient 2. Bottom: vertex-wise scatter plot (R = 0.63, p = 0.008, spin test). Vis: visual cortex. SM: somatomotor cortex. Aud: auditory cortex. MFG: middle frontal gyrus. IFG: inferior frontal gyrus. AG: angular gyrus. PreC: precuneus. MTC: middle temporal cortex. MeFC: medial frontal cortex. DMN: default mode network.

INF mode 1 also extended beyond the cortex to subcortical and cerebellar structures, with their first functional gradients aligning with either the cosine component (cerebellum: R = 0.87; thalamus: R = 0.95; brainstem: R = 0.92; striatum: R = 0.58; all p < 0.001, Moran spectral test) or the sine component (hippocampus: R = 0.87; amygdala: R = 0.75; all p ≤ 0.001, Moran spectral test), possibly depending on whether the dominant subcortical organization follows the cortical SA or VS axis (Figure 5). These results extend the observation beyond the cortex, demonstrating that the dominant functional gradients across cortical, subcortical, and cerebellar structures are unified as projections of a single, brain-wide intrinsic flow.

**Figure 5.**
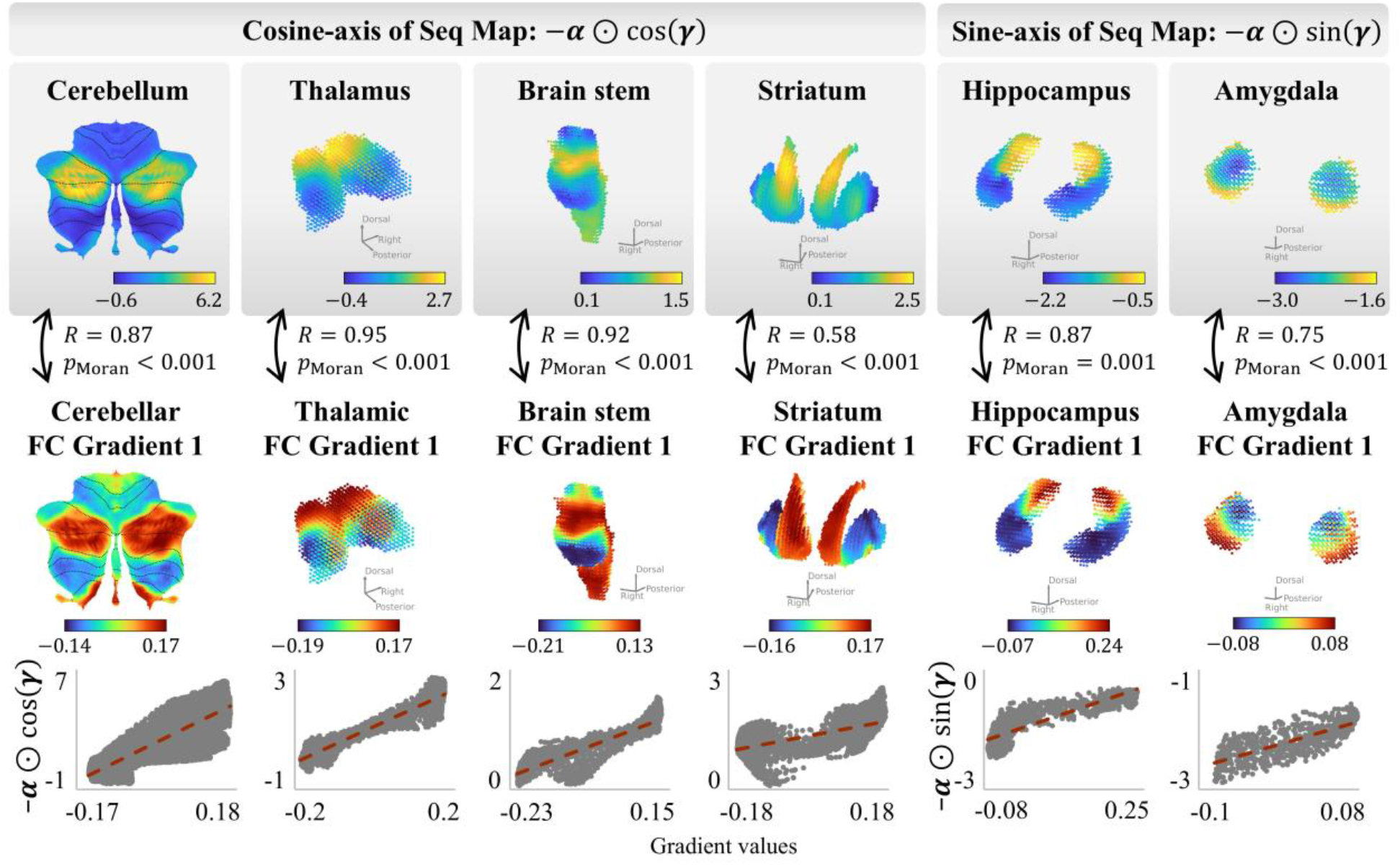
The principal flow extends across subcortical and cerebellar structures. The cosine-axis (−***α***_1_ ⊙ cos(***γ***_1_); left four columns) and sine-axis (−***α***_1_ ⊙ sin(***γ***_1_); right two columns) projections of the INF mode 1 flow sequence map are shown for the subcortical and cerebellar structures, each compared with its corresponding first FC gradient. Top row: flow sequence map projections. Middle row: FC gradient 1 maps. Bottom row: voxel-wise scatter plots with linear fits (red dashed lines). The cosine component aligns with the first FC gradient in structures whose dominant organization follows the cortical SA hierarchy (cerebellum: R = 0.87; thalamus: R = 0.95; brainstem: R = 0.92; striatum: R = 0.58; all p < 0.001), whereas the sine component aligns with the first FC gradient in structures whose organization follows the cortical VS axis (hippocampus: R = 0.87, p = 0.001; amygdala: R = 0.75, p < 0.001). Statistical significance was assessed using Moran spectral randomization (see Methods).

### INF 2–4: Higher-order flows capture lateralized network dynamics and triple networks

The next three modes revealed coordination structures among the brain’s major large-scale networks. INF mode 2 describes a lateralized flow between the left-dominant default mode network (DMN) and the central executive network (CEN) (Figure 6a; Supplementary Video 2), with its cosine component aligning with the DMN spatial map (posterior cingulate cortex seed-based FC; R = 0.64, p < 0.001, spin test), the CEN spatial map (R = −0.59, p < 0.001, spin test), and its left-hemispheric cosine component with the laterality index [33] (*R* = 0.89, *p* < 0.001, spin test; see Methods), reflecting left-hemispheric dominance (Figure 6b).

**Figure 6.**
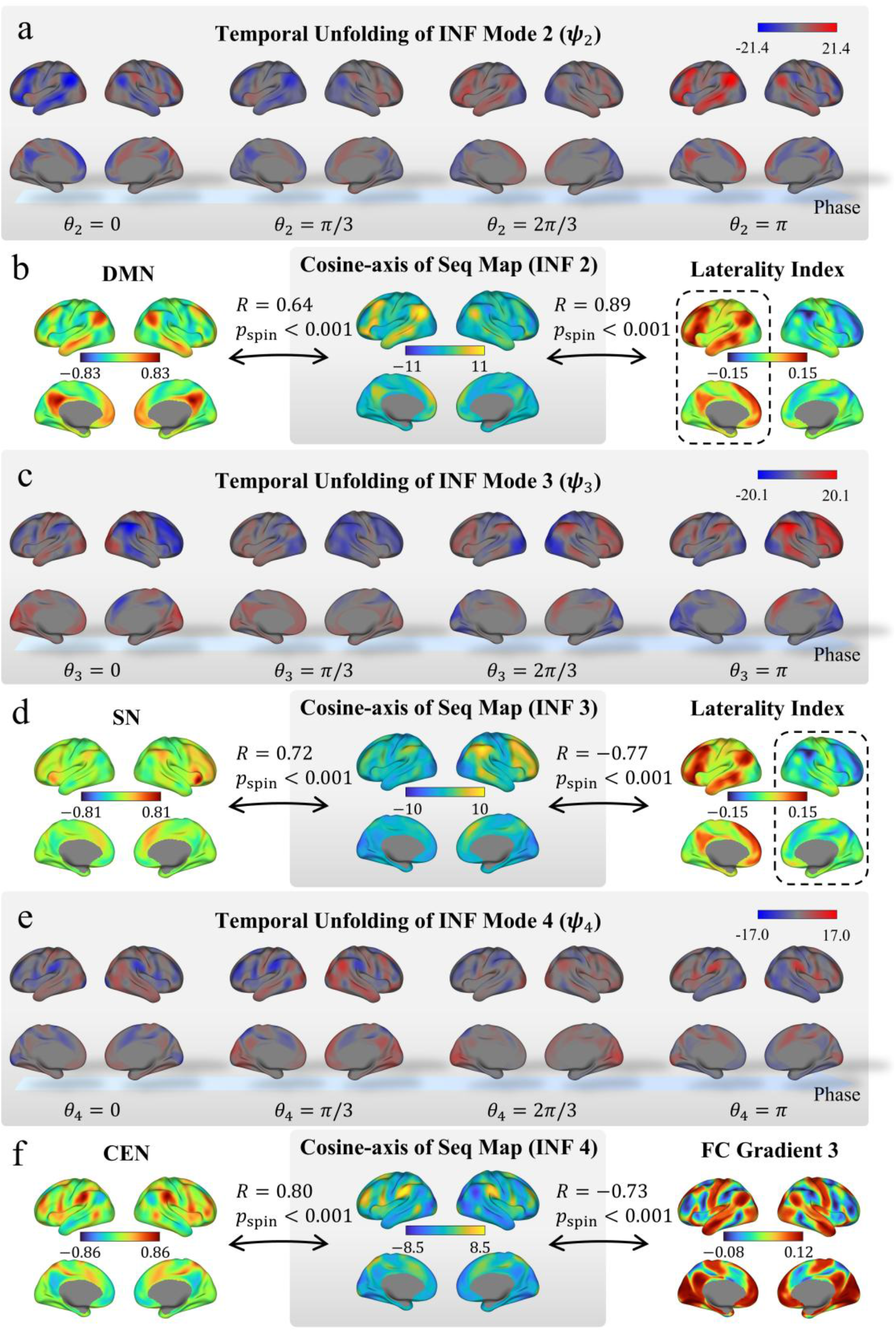
Higher-order INF modes capture lateralized network dynamics and triple network interactions. **(a)** Temporal unfolding of INF mode 2 (***ψ***_2_) across four phase values (*θ*_2_ = 0, π/3, 2π/3, π), showing a lateralized flow between DMN and CEN regions. **(b)** Spatial correspondence of the INF mode 2 cosine-axis of the flow sequence map with the DMN spatial map (posterior cingulate cortex seed-based FC; R = 0.64, p < 0.001) and the left-hemisphere component of laterality index (R = 0.89, p < 0.001), confirming left-hemispheric dominance. **(c)** Temporal unfolding of INF mode 3 (***ψ***_3_), showing a complementary flow between SN and representation regions. **(d)** Spatial correspondence of the INF mode 3 cosine-axis of the flow sequence map with the SN spatial map (right insula seed-based FC; R = 0.72, p < 0.001) and the right-hemisphere component of laterality index (R = −0.77, p < 0.001), confirming right-hemispheric dominance. **(e)** Temporal unfolding of INF mode 4 (***ψ***_4_), showing a flow along the representation–modulation (RM) axis (i.e., the third cortical FC gradient). **(f)** Spatial correspondence of the INF mode 4 cosine-axis of the flow sequence map with the CEN spatial map (supramarginal gyrus seed-based FC; R = 0.80, p < 0.001) and cortical FC gradient 3 (R = −0.73, p < 0.001). All spatial correlations are assessed using spin test with 1,000 permutations. Additional spatial correlations with CEN and RM-axis maps are reported in the main text. DMN: default mode network. SN: salience network. CEN: central executive network.

INF mode 3 exhibits a complementary pattern (Figure 6c; Supplementary Video 3): a flow between the right-dominant salience network (SN) and regions collectively referred to as the representation end of the representation–modulation (RM) axis (i.e., the third cortical FC gradient [34]—namely, DMN, visual, and SM systems). Its cosine component aligns with the SN spatial map (right insula seed-based FC; R = 0.72, p < 0.001, spin test), the third cortical FC gradient (R = −0.68, p < 0.001, spin test), and its right-hemispheric cosine component with the laterality index (R = −0.77, p < 0.001, spin test; Figure 6d). Together, INF modes 2 and 3 suggest that the functional lateralization of DMN [33, 35] and SN [36] is a dynamical property embedded within intrinsic flow, which are principal sources of laterality in brain dynamics.

INF mode 4 captures a flow along the RM axis (i.e., the third cortical FC gradient) [34], with one end anchored by the DMN, visual, and SM systems, and the other by CEN. Its cosine component strongly correlated with the CEN spatial map (supramarginal gyrus seed-based FC; *R* = 0.80, *p* < 0.001, spin test) and the third cortical FC gradient (*R* = −0.73, *p* < 0.001, spin test; Figure 6e, f; Supplementary Video 4).

Collectively, INF modes 2–4 reveal that the interactions among the triple networks, DMN, SN, and CEN [37], together with sensorimotor systems, are organized as distinct yet interrelated intrinsic flows. Notably, hemispheric lateralization embedded within their flow structure suggests that lateralization is an inherent operational property of triple network coordination. The remaining modes (5–13) capture additional coordination patterns and are presented in Supplementary Videos 5–13.

### Phase modulation reconstructs task activation while preserving intrinsic FC topology

A key argument of the INF framework is that diverse cognitive states emerge from phase modulation of intrinsic flows, without reconfiguring the flows themselves. To test the plausibility, we reconstructed task activation maps for 23 diverse cognitive tasks (multi-domain task battery (MDTB) dataset [38]; Figure 7a) using INF modes derived from HCP resting-state data. By modulating only the phases while flow amplitudes and modes are fixed (see Methods), we achieved high reconstruction accuracy of empirical task-evoked activation maps (mean spatial *R* = 0.73, max *R* = 0.88, min *R* = 0.64; Figure 7b), demonstrating that phase realignment of intrinsic flows is sufficient to generate the brain’s rich repertoire of task activation patterns. Crucially, this framework also explains the persistence of FC topology across cognitive states. If brain signals are superpositions of the same flows, correlation structure (i.e., FC topology) is determined by the INF modes themselves (***ψ***_*k*_) and their flow amplitude (*η*_*k*_), and is theoretically invariant to phase modulation, as phase differences integrate out over time (see Supplementary Result 3). Consistent with this, the theoretically reconstructed FC matrix aligned closely with both all task-state FC matrices (mean *R* = 0.78, max *R* = 0.83, min *R* = 0.68; Figure 7c) and the null condition FC matrix (*R* = 0.85; rest condition in MDTB; Supplementary Figure 8, top), including preservation of the anti-correlation structure between DMN and executive regions (Supplementary Figure 8, grey inset).

**Figure 7.**
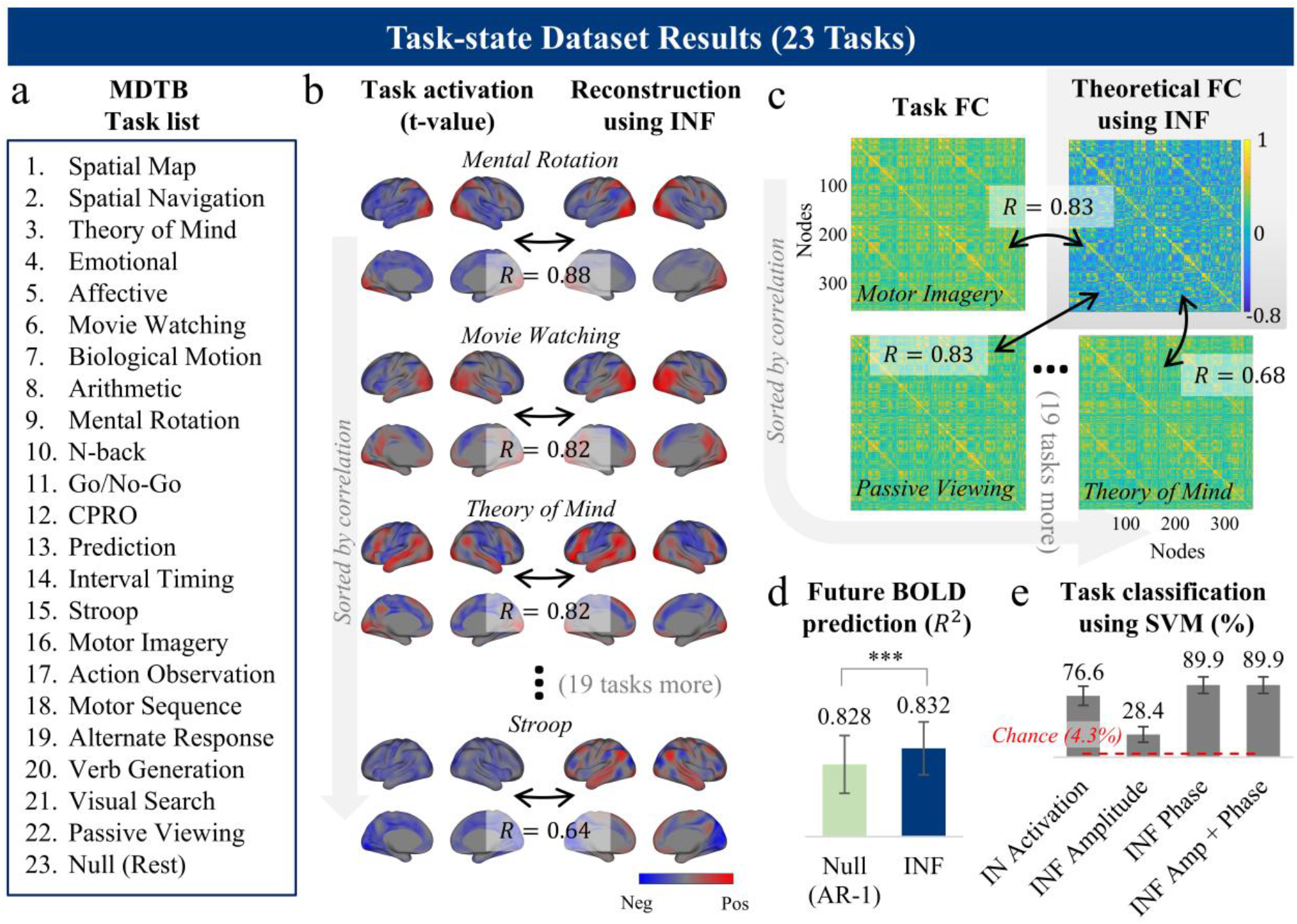
INF phase encodes cognitive states. **(a)** List of 23 task conditions from the multi-domain task battery (MDTB) dataset. **(b)** Task activation reconstruction using INF phase modulation. For each task, the empirical group-level activation map (t-value; left) is compared with the reconstruction obtained by modulating only the INF phases while holding INF modes and flow amplitudes fixed to those estimated from HCP resting-state data (right). Spatial correlations between empirical and reconstructed maps are shown (mean *R* = 0.73), and tasks are sorted by reconstruction accuracy. Full reconstruction results for all 23 task activation maps are provided in Supplementary Figures 7. **(c)** FC topology preservation. Left: empirical task-state FC matrices from MDTB data (Glasser 360 parcellation), sorted by correlation with the theoretically reconstructed FC. Right: FC matrix analytically derived from resting-state INF parameters (acquired from HCP), which is invariant to phase modulation (see Supplementary Result 3). Correlations between the theoretical FC and task-state FC matrices are shown (mean *R* = 0.78), sorted by correlation value (excluding null). Full reconstruction results for all task FC matrices are provided in Supplementary Figures 8. **(d)** Future BOLD signal prediction during MDTB task performance. INF modes derived from resting-state dataset (i.e. HCP) significantly outperformed the AR(1) null model in predicting future BOLD signals during MDTB task runs (*R*^2^ = 0.832 vs. 0.828; *p* = 1.4 × 10^-6^). Error bars denote confidence intervals **(e)** Task classification accuracy (leave-one-subject-out, linear SVM) across 23 MDTB tasks using four feature types: IN activation, INF amplitude, INF phase, and combined INF amplitude and phase. Red dashed line indicates chance level (4.3%). Error bars denote standard deviations.

Furthermore, prediction of future BOLD signals using resting-state-derived INF modes significantly outperformed the null AR(1) model during task performance (*R*^*2*^ = 0.832 versus *R*^*2*^ = 0.828, *p* = 1.4 × 10^-6^, *N* = 18, paired t-test; Figure 7d). Together with the preservation of FC topology across task states, these results indicate that the intrinsic flow structures are largely preserved across different cognitive states.

### INF phase encodes cognitive states; Amplitude is trait-like

Lastly, we examined the information that phase encodes. If cognitive states are realized through phase dynamics, then phase should carry task-relevant information. Consistent with this prediction, INF phases not only differentiated ongoing task conditions but also significantly outperformed both flow amplitudes and intrinsic network activation levels in decoding cognitive states.

In a leave-one-subject-out (LOSO) classification of 23 MDTB tasks using a linear support vector machine (SVM), models trained on INF phase features achieved 89.9 ± 10.3% accuracy (mean ± standard deviation), compared to 76.6 ± 12.0% for IN activation-based features and 28.4 ± 9.8% for INF amplitude features (Figure 7e). Combining phase and amplitude features yielded no improvement. To further test this finding, we applied the same approach to an audiovisual speech perception dataset comprising four conditions sharing a common task goal (speech perception) but differing in sensory modality (audiovisual, visual only, audio only, and rest [39]; Supplementary Figure 9a), in which activation maps were highly similar across conditions (e.g., *R* = 0.72 between audio-only and audiovisual stimuli; Supplementary Figure 9b). Phase-based classification achieved 95.0 ± 11.1% accuracy, outperforming amplitude-based and IN activation-based classification (46.3 ± 22.5% and 87.9 ± 17.5%, respectively; Supplementary Figure 9c).

Despite their limited relevance for state discrimination, flow amplitudes exhibited significant heritability, as estimated by the additive genetic, common environmental, and unique environmental (ACE) modeling [40] using HCP kinship data (*h*^2^ = 0.41, 95% CI: [0.32, 0.48], *p* = 0.0002, *N* = 945). Moreover, four of the 13 modes showed consistent positive correlation between amplitudes and general cognitive ability [41] estimated from HCP behavioral data (false discovery rate (FDR) < 0.05; Supplementary Figure 10b).

Taken together, the results suggest a conceptual dissociation: INF phase operates as a flexible state variable that configures moment-to-moment cognitive dynamics, whereas flow amplitude functions as a trait-like parameter that likely reflects individual differences in cognitive capacity.

## Discussion

Neuroscience has long been guided by the principle of functional segregation—that distinct brain regions serve distinct computational roles [42-44]—which naturally frames cognition as a matter of recruiting appropriate regions and reconfiguring their interactions. Under this view, flexible cognition requires flexible architecture: the system must reorganize itself (e.g., recruiting different regions, altering connectivity, or switching network configurations) to meet each new demand. Our work offers an alternative perspective. The INF framework demonstrates that the brain need not alter the coordination structure underlying its functional architecture to generate diverse cognitive states. Instead, what may appear like recruitment of different regions or reconfiguration of network interactions can be parsimoniously accounted for by shifts in the relative timing (i.e., phase dynamics) of stable intrinsic flows, whose amplitudes and sequential structure remain preserved, thus reconciling flexibility with stability. In other words, flexible cognition may not require flexible architecture, but rather the retiming of a fixed one—offering a resolution to the long-standing stability–flexibility duality.

Within this framework, the paradoxical stability of FC topology across different cognitive states becomes a natural consequence of having consistent flows: temporal correlations are invariant under phase modulation, so retiming the flows leaves their correlational structure unchanged. Likewise, the organizational features that the flows generate (e.g., functional gradients, canonical networks) are preserved across tasks. This perspective also accounts for the widely observed spatial similarity between resting-state and task-state brain activity patterns [5, 20, 45]: both reflect snapshots of the same flows. This situates resting-state and task-state dynamics within a single descriptive level: they are not separate phenomena but different manifestations of the same underlying coordination structure.

Situating such stable flows as the fundamental units of large-scale brain organization necessitates a revision of how task-evoked neural activity is interpreted. In conventional accounts, task-evoked deactivation is understood as suppression or disengagement of specific regions [46-50]. Within the INF framework, however, deactivation reflects negatively constructive interference among concurrent flows at particular phase alignments; and activation, conversely, reflects positively constructive interference. The finding that deactivation deepens with increasing task difficulty [51-53] follows naturally from this view: greater cognitive demand produces more sustained phase alignment, resulting in sharper interference rather than stronger suppression of individual regions. Because the flows themselves persist, all brain regions remain continuously active [54, 55] (see also [56]); what changes across cognitive states is not which regions are engaged, but how their ongoing dynamics align over time.

These findings challenge the prevailing assumption in systems neuroscience that regional activation amplitude is the primary variable for characterizing cognitive states [15] and modeling cognitive control [16, 17]. In the INF framework, tasks producing nearly identical activation maps can nevertheless occupy distinct positions in phase space. This was supported by the audiovisual speech perception paradigm, in which activation maps were highly similar across conditions, yet phase-based classification achieved near-perfect accuracy (95.0%). Moreover, across 23 diverse tasks, INF phases consistently outperformed both network activation levels and flow amplitudes in decoding cognitive states, despite the INF bases being derived from resting-state coordination structure. We note that phase is estimated from the same task activation time courses and therefore cannot carry information beyond them. Thus, the relevant point is the sufficiency of timing alone to characterize cognitive states, even for tasks with similar activation maps. This calls into question the intuitive assumption that cognitive adaptation is achieved primarily by recruiting the appropriate brain regions (or initiating network flows), an assumption that also leaves the stability discussed above unexplained. Instead, these results collectively suggest that cognitive states are carried by *when* intrinsic flows align in time (i.e., phase) rather than by *how strongly* they are expressed, pointing toward a reconceptualization of both brain state representation and the control variables that govern flexible cognition (see also [57-59]).

Our framework unifies features of functional organization that have often been studied in isolation. Notably, the principal flow alone recapitulates the dominant functional gradients across cortex, cerebellum, and subcortex, suggesting that region-specific gradient analyses [32, 34, 60, 61] reflect projections of a single brain-wide coordination structure. The two dominant cortical gradients—SA and VS axes—have been treated as independent dimensions, and while the SA axis has been linked to subcortical and cerebellar organizations, the VS axis has remained less clear [34]. Our results indicate that both can be understood as projections of a single flow, with the VS axis capturing the timing difference between visual and SM systems during their interaction with association cortex (e.g., DMN)—potentially related to their distinct hierarchical depth in processing [62]. Moreover, by embedding cortical and subcortical/cerebellar structures within a unified flow, this framework offers a unique opportunity to understand subcortical and cerebellar contributions to large-scale information integration. That the third cortical gradient corresponds to a distinct flow (INF mode 4) further supports the view that FC gradients broadly reflect static projections of the brain’s major intrinsic flows. By situating these intrinsic organizational features within temporally structured flows that underlie cognitive diversity, the framework provides a principled bridge between resting-state functional architecture and task-evoked cognitive dynamics.

The INF framework shares a conceptual foundation with recent eigenmode-based approaches, such as geometric eigenmodes [63] and connectome harmonics [64], in decomposing whole-brain activity into superpositions of large-scale basis patterns. However, these approaches derive their bases from structural geometry without explicitly modeling temporal structure, and justify linearity through linearization near a fixed point [65, 66] or Fourier theory [64]. The INF framework differs on both counts: it embeds temporal order (i.e., flow sequence, capturing both past and future evolution) within the basis itself, and is based on Koopman operator theory, which establishes that nonlinear dynamics can be represented linearly when embedded in a sufficiently rich observable space [67, 68]. This theoretical grounding motivates a key methodological choice: modeling group-level dynamics at full vertex/voxel resolution without prior parcellation, so that high-dimensional fMRI signals serve as the expansive observable basis that Koopman theory requires, while multi-subject estimation increases the effective time-series length needed for convergence toward the true Koopman modes [69, 70]. We have shown that the phase information emerging from this temporally structured decomposition is the crucial carrier of cognitive state information, which cannot be yielded by other eigenmode decompositions.

Finally, the INF framework provides a principled decomposition of brain dynamics into group-level coordination structures (i.e., INF modes), subject-specific fingerprints (e.g., spatial maps of INs, amplitudes of flows; see Methods), and residual fluctuations (see Methods), enabling direct between-subject comparison of dynamical properties within common coordinates. Moreover, within this decomposition, the amplitude–phase dissociation conceptually separates trait and state contributions [71, 72]: flow amplitudes capture trait-level individual differences (i.e., heritable, stable across cognitive states, and predictive of general cognitive ability), while INF phases encode instant cognitive state changes. Whether pathological reductions in flow amplitude, or qualitative changes in flow modes themselves, characterize clinical populations remains an important direction for future work.

In summary, the INF framework demonstrates that the brain’s diverse cognitive repertoire can emerge from phase dynamics over a fixed set of intrinsic flows without reconfiguring its architecture. By grounding this principle in a formal decomposition with empirically testable predictions, the framework offers a unified language for linking resting-state organization, task-evoked activation, and individual differences within a single descriptive level. We anticipate that this perspective will open new avenues for understanding how large-scale neural coordination gives rise to flexible cognition in health and disease.

## Methods

### HCP dataset and preprocessing

We employed resting-state fMRI data from the HCP S1200 release [73]. Data collection was approved by the Institutional Review Board at Washington University in St. Louis, and all participants gave informed consent. Our data use strictly adhered to the HCP Data Use Agreement.

Each participant completed four 15-minute resting-state fMRI runs over two separate days. On each day, two runs were acquired with opposing phase-encoding directions—left-to-right (L/R) and right-to-left (R/L)—while participants kept their eyes open and fixated. Detailed acquisition procedures are described in previous work [74].

For this study, we included participants who completed either one day of resting-state scan (i.e., the REST1 or REST2 session), which comprised both L/R and R/L scans. Participants were excluded if they had a mini-mental state examination (MMSE) score of 26 or lower, which may suggest cognitive impairment in this young adult sample [75]. We also excluded scans in which the average root-mean-squared frame-to-frame head motion (as recorded in *Movement_Relative_RMS_mean*.*txt*) exceeded 0.15 mm in either direction [76, 77]. Additionally, participants were excluded if their R/L scan data exhibited preprocessing errors, as documented in the HCP public resource “HCP-Young Adult Data Release Updates: Known Issues and Planned Fixes” (https://wiki.humanconnectome.org/). Then, the REST1 sample consisted of 951 participants (female = 520, mean age = 28.71, std = 3.69), and the REST2 sample consisted of 884 participants (female = 484, mean age = 28.63, std = 3.72).

For the assessment of BOLD signal prediction, we retain only genetically unrelated subjects from REST1 cohorts to avoid inflating reproducibility or predictive accuracy through genetic similarity (*N* = 420, female = 231, mean age = 28.79, std = 3.68). Then, we first drew a discovery dataset of 210 individuals, randomly splitting them into two equal groups (*N* = 105 each; including 55 and 59 females respectively): one group served as the training dataset for estimating group-level coordination structures (discovery-training set), and the other as the test dataset for evaluating prediction performance (discovery-test set). To assess reproducibility, we then independently drew a replication dataset of another 210 participants and applied the identical split (replication-training and replication-test sets; *N* = 105 each; including 61 and 56 females respectively).

All MRI scans were acquired at Washington University in St. Louis using a customized Siemens 3T Skyra scanner with a multiband echo-planar imaging (EPI) sequence for accelerated imaging. Structural images were obtained at 0.7 mm isotropic resolution, while resting-state fMRI scans had 2 mm isotropic resolution and a TR of 0.72 seconds. Full details of data acquisition and preprocessing are available in prior studies [73, 74].

We used the 2017 release of preprocessed surface-based CIFTI-format fMRI data, which had undergone a sulcal depth-based surface registration (MSMSulc) and ICA-FIX denoising to remove structured artifacts [74]. Spatial smoothing (6 mm FWHM Gaussian kernel) was applied to both surface and volume components using the Connectome Workbench [78]. No temporal filtering or nuisance regression was applied. Prior to further analysis, each dataset was temporally demeaned and variance-normalized (i.e., z-scored across time for each vertex and voxel).

### MDTB dataset and preprocessing

We used the publicly available multi-domain task battery (MDTB) dataset [38] (OpenNeuro ds002105). This dataset comprises task fMRI data from 24 right-handed individuals (16 female, 8 male; mean age: 23.8, std: 2.6 years) collected at Western University under an experimental protocol approved by the institutional review board. All data were acquired on a 3T Siemens Prisma scanner at Western University using a multiband EPI sequence (TR = 1 s; voxel size = 2.5 × 2.5 × 3.0 mm^3^; 48 slices). Gradient echo fieldmaps were acquired for distortion correction, and a T1-weighted anatomical scan was collected at 1 mm isotropic resolution. Participants performed 26 distinct cognitive tasks spanning motor, cognitive, affective, and social domains across two scanning sessions, each consisting of two imaging runs. Sessions were separated by either 2–3 weeks or 1 year across participants. Tasks were performed in 35-second blocks (5-second instruction screen followed by 30 seconds of continuous performance), with all tasks within a session completed in a single imaging run to ensure a common baseline. Of the 24 participants, resting-state fMRI was additionally collected for 18 individuals. For this study, we treated the two N-back variants (verbal and picture) as a single task condition and similarly combined three movie-watching conditions (natural, landscape, and romance) into a single task, yielding 23 task conditions for analysis. Full details of the experimental tasks and conditions are provided in the original publication [38].

Preprocessing was performed using fMRIPrep 25.2.3 [79], which included head-motion correction, fieldmap correction, spatial normalization using T1-weighted structural images, and surface registration to produce CIFTI-format grayordinate files; no slice-timing correction was applied (see Supplementary Method 1 for full preprocessing details provided by fMRIPrep software). Spatial smoothing was performed on the cortical surface and subcortical volumes using Connectome Workbench [78] with a Gaussian kernel of 8 mm FWHM (chosen to approximately match the effective smoothness of the HCP data, which were acquired with 2 mm prior smoothing and further smoothed at 6 mm FWHM in this work). Additional denoising consisted of regressing out 24 motion parameters, mean white-matter and CSF time series, and linear and quadratic trends, followed by bandpass filtering at 0.008–0.15 Hz. After denoising, the data was temporally demeaned and variance normalized.

### Audiovisual speech dataset and preprocessing

We used the publicly available audiovisual speech perception dataset [39] (OpenNeuro ds003717). The dataset comprises 60 right-handed participants (45 female; mean age 22.4 ± 3.2 years), all native speakers of American English with self-reported normal hearing. All participants provided informed consent under a protocol approved by the institutional review board at Washington University in St. Louis. Data were acquired on a 3T Siemens Prisma scanner at Washington University in St. Louis using a multiband EPI sequence (voxel size = 2.0 mm isotropic; sparse imaging design with TR = 2.47s and acquisition time = 0.77s). A T1-weighted MPRAGE structural scan was collected at 0.8 mm isotropic resolution. Participants performed a speech perception task across seven conditions: audiovisual speech at four signal-to-noise ratios (quiet, +15, 0, −5, −10 dB SNR), visual-only (lipreading), and auditory-only. Stimuli consisted of 1.5-second video recordings of a female actor speaking single words, presented in blocks of five experimental trials plus two null trials. Five perception runs (∼5.5 minutes each) were completed per session. Full details of the experimental tasks and conditions are provided in the original publication. For this study, we considered four conditions: audiovisual speech in quiet, visual-only, auditory-only, and null trials (rest).

Preprocessing was performed using fMRIPrep 25.2.3 [79], which included head-motion correction, spatial normalization using T1-weighted structural images, and surface registration to produce CIFTI-format grayordinate files; no slice-timing correction was applied (see Supplementary Method 2 for full preprocessing details). Spatial smoothing, additional denoising, and variance normalization were identical to the MDTB dataset described above.

### INF mode estimation

The INF mode estimation procedure consists of two main stages: (1) IN discovery and (2) dynamic coherence modeling. A schematic overview of the estimation is provided at STEP 1 and 2 in Figure 2.

The IN discovery is performed using group independent component analysis (GICA) [80]. To this end, we first conducted group-level principal component analysis (PCA) using MELODIC’s incremental group-PCA (MIGP) [81]. The data were then reduced to the target dimensionality *Q*, corresponding to the number of INs to be extracted. The reduced data were whitened to ensure unit variance and zero covariance across components. Next, the Infomax ICA algorithm implemented in the GIFT toolbox (http://trendscenter.org/software/gift) was applied to the reduced and whitened dataset, yielding a matrix of group-level IN maps, denoted as *S*, whose columns correspond to different INs. Using the IN map matrix, we extracted IN time-course for all subjects using dual regression [82-84]:

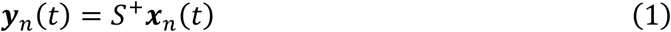

where ***x***_*n*_(*t*) ∈ ℝ^*V*^ and ***y***_*n*_(*t*) ∈ ℝ^*Q*^ represent participant’s voxel-level fMRI activation and the IN activation vectors at time *t* for subject *n. S*^+^ denotes the pseudo-inverse of *S*, and *V* is the number of voxels. Noise-component removal at this stage [85, 86], when needed, can improve the robustness of the subsequent procedure [87]. In the present study, we did not perform additional noise removal because the data had already been denoised using ICA-FIX.

For dynamic coherence modeling, we first modeled the IN activity as a group-level linear system [88]:

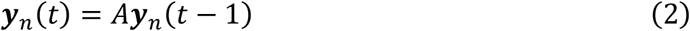

where *A* ∈ ℝ^*Q*×*Q*^ is the group-level transition matrix capturing interactions between INs. The matrix *A* can be estimated via least squares by minimizing prediction error across all subjects and all time points [88]:

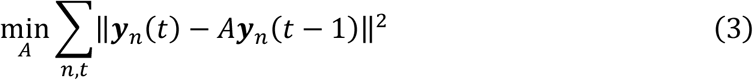

To reduce systematic estimation bias, we adopted the forward–backward dynamic mode decomposition (fbDMD) algorithm [87], as described in the prior work [28] for estimating *A*.

After estimating *A*, we performed eigenvalue decomposition to extract the group-level INF modes. The eigenvectors ***ψ***_*k*_ ∈ ℂ^*Q*^ and their corresponding eigenvalues *λ*_*k*_ ∈ ℂ satisfy:

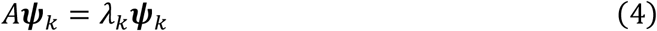

where *k* is eigenvalue index. Each eigenvalue 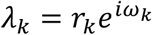 corresponds to the intrinsic evolution coefficient *d*_*k*_ described in the framework, governing the spontaneous temporal evolution of each INF mode: *r*_*k*_ determines the persistence of the flow amplitude over time, and *ω*_*k*_ determines the speed at which the phase advances per time step.

Since *A* is a real value matrix, its complex eigenvalues and eigenvectors appear in conjugate pairs. In this work, an INF mode is defined as a pair of complex-conjugate eigenvectors, which jointly represent coherent and directional signal propagation across networks (INs). Real-valued eigenvectors, although mathematically valid solutions of Eq. (4), do not encode flow-like dynamics and are therefore not considered INF modes. As a result, while *A* has *Q* eigenvectors in total, only the complex-conjugate pairs are retained for INF mode analysis, yielding at most *K* ≤ *Q*/2 INF modes.

### Fingerprinting after mode estimation

To relate INF modes to individual brain activity, we derive spatiotemporal fingerprints in two steps: spatial fingerprint and temporal fingerprinting (STEP 3 and 4 in Figure 2). For spatial fingerprinting, we extracted the subject-specific IN spatial maps (*S*_*n*_ ∈ ℝ^*V*×*Q*^) using dual regression [82]. Then, the subject-specific voxel-level expression of each INF mode (i.e., *spatial fingerprints*) was obtained by multiplying the subject-specific IN spatial maps with the corresponding INF mode vector:

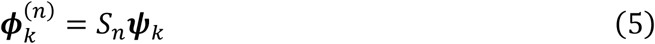

For temporal fingerprinting, each subject’s voxel-level BOLD activity ***x***_*n*_(*t*) was represented by the corresponding subject’s spatial fingerprints 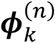:

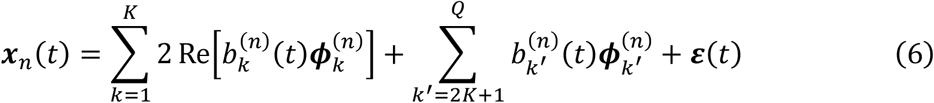

Here, 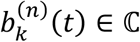, denotes the flow coefficient corresponding to the *k*-th INF mode, reflecting the activation at time *t*. The terms indexed by *k*^′^ correspond to real-valued eigenvectors of **A**, which are treated as non-propagating components. ***ε***(*t*) represents residual activity.

The flow coefficients were estimated by projecting each voxel-level BOLD activity vector onto the subspace spanned by the voxel-level expressions of the eigenvectors of **A**. Specifically, for subject *n*, we constructed a voxel-level basis matrix

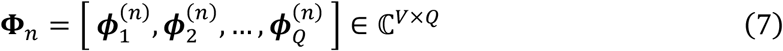

where each column 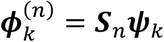 represents the voxel-level expression of the *k* -th eigenvector, including both complex-conjugate INF mode vectors and real-valued eigenvectors. Given the BOLD activity vector ***x***_*n*_(*t*) ∈ ℝ^*V*^, the corresponding flow coefficient vector ***b***_*n*_(*t*) ∈ ℂ^*Q*^was obtained via a least-squares projection:

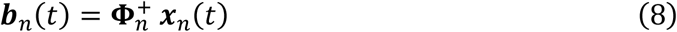

where + denotes the Moore–Penrose pseudoinverse. This projection yields complex-valued coefficients for INF modes and real-valued coefficients for non-propagating (real-valued) modes. The estimated coefficients were then used to reconstruct the BOLD activity as described in Eq. (6).

The one-step-ahead predicted BOLD signal is then modeled as:

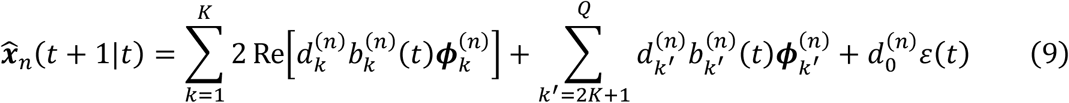

Here, 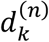 captures the temporal evolution dynamics of each INF mode. An additional coefficient 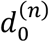 is included to model residual autocorrelation. The evolution coefficients 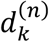 were estimated to optimally match the predicted signal 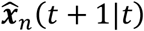 with the observed signal ***x***_*n*_(*t* + 1):

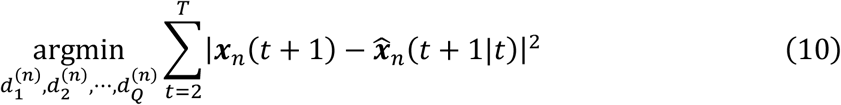

A least-square solution for such evolution coefficients estimation is provided in ref [28], Supplementary Method 2.

Then, using estimated 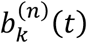 and 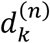, three metrics were computed for each INF mode (i.e., *temporal fingerprints*): for *k*-th INF mode, (1) average amplitude 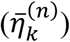, (2) persistence rate 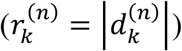, and (3) progression speed 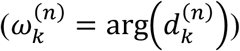. The average amplitude quantifies how consistently the corresponding INF mode is engaged during the scan and is defined as the temporal average of its amplitude:

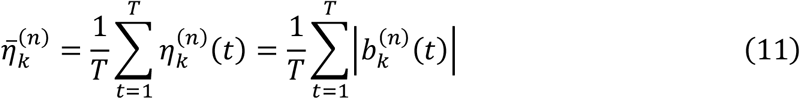

In this way, the INF framework describes a formal generative relationship, demonstrating how a common coordination blueprint (i.e., INF modes) is uniquely instantiated through subject-specific parameters (i.e., spatial and temporal fingerprints) to give rise to the diverse complexity of observable brain dynamics.

### BOLD signal prediction using INF modes

For each subject in the test set, the resting-state fMRI scan was divided into two distinct segments. The test-fitting segment (first 60% of the time series, ∼9 minutes) was used to spatiotemporal fingerprinting. The test-validation segment (remaining 40%, ∼6 minutes) was held out for evaluating the predictive performance.

Prediction accuracy was evaluated using the test-validation segment, with the error at time *t* defined as:

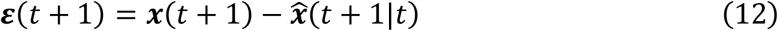

To quantify predictive performance across different brain regions, we used *R*^2^ which measures the proportion of variance explained by the model. Because BOLD signal variance differs across brain locations, *R*^2^ was computed per vertex/voxel and then averaged across all vertex/voxel:

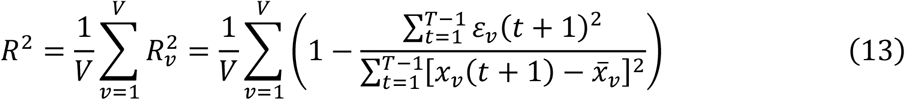

Here, *x*_*v*_(*t* + 1) denotes the BOLD signal at vertex/voxel *v* and time *t* + 1; *ε*_*v*_(*t* + 1) is the corresponding prediction error; and 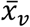 is the mean activation over time: 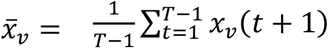.

### Null model

To assess the predictive utility of the coordination structures, we compared their performance to a baseline model. We adopted a first-order autoregressive AR(1) null model, which captures only the temporal autocorrelation between consecutive time points:

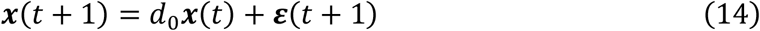

Here, *d*_0_ is a scalar coefficient representing the lag-1 autocorrelation, and ***ε***(*t*) denotes a white noise error term. The coefficient *d*_0_ was estimated analytically using the following closed-form expression:

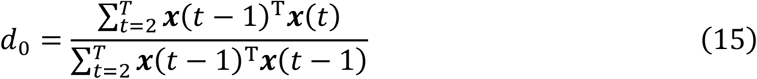

where ***x***(*t* − 1)^T^ indicates the transpose of ***x***(*t* − 1).

### Canonical features in resting-state fMRI

#### Functional connectivity (FC) gradients

To obtain canonical features of resting-state fMRI, the spatially smoothed HCP S1200 data were further band-pass filtered in the frequency range of 0.01–0.1 Hz. Cortical FC gradients were computed by applying diffusion embedding to subject-averaged vertex-to-vertex FC matrices using the BrainSpace toolbox [89]. For each subject, FC matrices were constructed by computing Pearson’s correlation coefficients between all pairs of cortical vertices. The resulting FC matrices were Fisher’s z-transformed (tanh^−1^), averaged across subjects, and then inverse Fisher-transformed (tanh) to obtain the group-level FC matrix. Diffusion embedding was performed using the default parameters in the BrainSpace toolbox (kernel: normalized angle; sparsity: 0.9; alpha: 0.5; automatic estimation of diffusion time).

To compute FC gradients for subcortical structures and the cerebellum, we generated FC profiles between these regions and the cerebral cortex. Specifically, for each subcortical region (hippocampus, thalamus, amygdala, brainstem, and striatum) and the cerebellum, group-level FC matrices were calculated between all voxels in the corresponding subcortical/cerebellar region and all cortical vertices, yielding one FC matrix per region. These matrices, representing voxel-wise cortical connectivity patterns, were used as input for diffusion embedding via the BrainSpace toolbox. Similarity between voxels was quantified using a normalized angle kernel, ensuring that voxels with similar cortical connectivity profiles were positioned closely in the resulting gradient space. The embedding was performed using the same parameters as the cortical gradient analysis.

#### Seed-based functional connectivity (FC)

FC was computed from cortical vertex time series after applying global signal regression (GSR). For GSR, the global mean time series—obtained by averaging signals across all cortical vertices—was regressed out from each vertex’s time series. The resulting residuals were then used to estimate FC, defined as the Pearson correlation between the residual time series of each seed ROI and those of all other cortical vertices.

#### Laterality index in vertex resolution

We quantified hemispheric lateralization using the mean laterality index (MLI) originally defined by ref [33] and detailed in Supplementary Method 5.11 of ref [28]. In the present study, the computation was identical to the ROI-based procedure described therein, except that calculations were performed for each cortical vertices rather than for pre-defined ROIs.

Briefly, the MLI represents the average hemispheric bias of a signal’s synchronization over the entire scan, with positive values indicating stronger coupling to the left hemisphere and negative values indicating stronger coupling to the right hemisphere. This is achieved by first computing the dynamic laterality index (DLI) within sliding time windows as the difference between correlation with the left-hemisphere global signal and correlation with the right-hemisphere global signal, and then averaging these DLI values across all windows.

### Task-evoked activation and FC analysis (MDTB and audiovisual perception)

For the MDTB dataset, task activation maps were estimated using a general linear model (GLM) on the preprocessed CIFTI grayordinate data (yet to be additionally denoised). For each imaging run, each of the 23 task conditions was modeled as a separate regressor (excluding the instruction period), represented as a boxcar function convolved with the SPM canonical hemodynamic response function. The GLM additionally included 12 motion-related nuisance regressors (six rigid-body motion parameters and their temporal derivatives). The GLM was estimated using SPM’s restricted maximum likelihood (ReML) function with an AR(1) covariance basis [90], and a high-pass filter at 128s was applied. Activation estimates for each subject and task condition were obtained by averaging the corresponding beta coefficients across runs. Group-level activation maps were then obtained by performing one-sample t-tests (*N* = 24) on the subject-wise beta maps for each task condition, yielding a t-value map per condition.

FC was calculated from the preprocessed and denoised CIFTI data parcellated using the Glasser cortical atlas (360 ROIs) [91]. To obtain condition-specific FC matrices for each of the 23 task conditions, Pearson correlations between ROI time series were computed using the HRF-convolved task-condition regressor as a weighting function, with negative values set to zero to ensure non-negative weights. This yielded one 360 × 360 FC matrix per condition per run for each subject. FC matrices were first Fisher r-to-z transformed, and then matrices from the same condition were averaged across runs within each subject. These within-subject averages were subsequently averaged across all subjects and converted back to correlation coefficients using the inverse Fisher transformation (tanh).

For the audiovisual speech perception dataset, task-evoked activation maps were estimated using the same GLM procedure. All eight experimental conditions were modeled in the GLM with a uniform trial duration of 2.3s to ensure consistency and prevent overlap with subsequent trial onsets; four conditions (audiovisual speech in quiet, visual-only, auditory-only, and rest) were retained for subsequent analyses.

### Task-evoked activation reconstruction using INF phase

For each of the 23 task conditions, the group-level activation map (t-value map, ***T*** ∈ ℝ^*V*^) was projected onto the IN spatial maps (*S* ∈ ℝ^*V*×*Q*^) using dual regression, yielding IN loading coefficients (***y*** = *S*^+^***T***). These loadings were subsequently projected onto the INF mode vector space to obtain flow coefficients *b*_*k*_ for each mode *k*. To isolate the contribution of phase information, each coefficient was expressed in polar form, 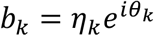. The amplitude |*b*_*k*_| was then replaced with the group-averaged INF amplitude 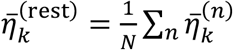 estimated from the HCP REST1 dataset, resulting in modified coefficients 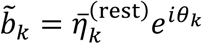. Thus, we only use phase information during reconstruction. Then, the reconstructed network activation vector was computed as 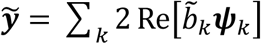 where ***ψ***_*k*_ denotes the INF mode vector from HCP dataset (see “INF mode estimation”). Finally, the full-resolution reconstructed activation map was obtained by projecting 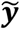 back through the group-level IN spatial maps *S*,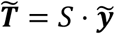.

### FC matrix reconstruction using INF modes

FC matrices were analytically reconstructed using INF modes and group-averaged INF amplitudes estimated from the HCP REST1 dataset. For each mode *k*, the scaled voxel-level mode was first obtained by projecting the INF vector ***ψ***_*k*_ onto the group-level IN spatial maps *S*, then scaled by the group-average INF amplitude 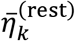 from HCP REST1 dataset:

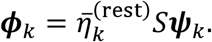

The resulting voxel-level mode ***ϕ***_*k*_ ∈ ℂ^*V*^ (*V* : number of voxels) was then parcellated according to the Glasser atlas (360 ROIs), yielding 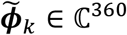. For each region *i*, we extracted the magnitude and argument of 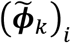, defining 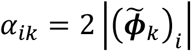 and 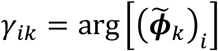, where the factor of 2 accounts for the contribution of the complex-conjugate mode pair.

When multiple INF modes contribute to brain activity, the Pearson correlation between regions *i* and *j*, under long-time averaging, reduces to a closed-form expression due to the orthogonality of distinct oscillation frequencies (see Supplementary Result 3 for derivation):

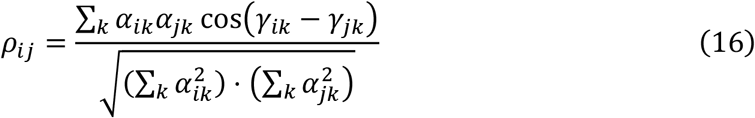

### Task-evoked INF amplitude/phase and IN activation extraction for classification

To extract task-evoked INF features, flow coefficient time courses *b*_*k*_(*t*) were estimated for each mode *k*. In principle, this can be achieved by projecting IN time courses ***y***(*t*) ∈ ℝ^*Q*^ onto the INF mode vectors (see eq. 8). In this analysis, we adapted the optimal amplitude estimation approach of ref [92] using the denoised voxel time series ***x***(*t*) ∈ ℝ^*V*^ and voxel-level INF modes constructed from group-level IN spatial maps (*S* ⋅ ***ψ***_*k*_). We used this approach to incorporate adjacent time points (***x***(*t* − 1), ***x***(*t*), ***x***(*t* + 1)) when estimating *b*_*k*_(*t*), leveraging the temporal evolution information inherent in the flow structure and yielding more stable flow coefficient time courses.

Then, each complex-valued flow coefficient *b*_*k*_(*t*) was decomposed into its magnitude *η*_*k*_(*t*) and argument *θ*_*k*_(*t*), yielding INF amplitude and INF phase time courses, respectively. Because the phase *θ*_*k*_(*t*) is a circular variable on [−*π, π*], it was further decomposed into its cosine and sine components, cos(*θ*_*k*_(*t*)) and sin(*θ*_*k*_(*t*)), to enable standard linear modeling.

Task-evoked estimates for each feature type were obtained using a GLM with the same HRF-convolved task-condition regressors used for the activation analysis. Since the input time courses were already denoised, no additional motion regressors or bandpass filtering were applied. Temporal autocorrelation was accounted for using an AR(1) covariance basis set, estimated via ReML. The GLM was performed separately for each of the three feature types— amplitude, cos(*θ*_*k*_(*t*)), and sin(*θ*_*k*_(*t*)) —and for each imaging run. Beta estimates for corresponding task conditions were averaged across runs, yielding one feature vector per condition per subject for classification (see Supplementary Figure 6 for schematic overview).

For comparison, IN time courses ***y***(*t*) were used to the same GLM procedure to obtain task-evoked IN activation beta values, which served as activation-based features for classification.

### Future BOLD prediction for MDTB dataset

Of the 24 participants from MDTB dataset, 18 also completed independent resting-state sessions (other than the null condition). Future BOLD prediction was performed for these 18 individuals. Using the INF modes estimated from the HCP REST1 dataset (*N* = 951), spatiotemporal fingerprinting was performed on these MDTB resting-state runs (see “Fingerprinting after mode estimation”), and future BOLD signal prediction accuracy for the preprocessed and denoised task session data was evaluated following the same procedure as for the HCP dataset (see “BOLD signal prediction using INF modes”).

### Heritability analysis

To evaluate the heritability of spatiotemporal fingerprints, we leveraged twin data obtained from the HCP. Individuals who were missing zygosity data or key covariates—such as age, sex, and structural measures—were removed from the analysis. The final sample comprised 945 participants: 91 monozygotic (MZ) twin pairs, 91 dizygotic (DZ) twin pairs, and 120 singletons. Using zygosity information, we fitted an ACE model that decomposes the total phenotypic variance into three components: genetic factors (A, representing heritability), shared environmental (C), and unique environmental factors (E). We employed the accelerated permutation inference for ACE models (APACE) [40], with each participant’s mean INF amplitude as the input phenotype. Prior to analysis, several nuisance variables were regressed out, including age, age squared, sex, the interaction between age and sex, the interaction between sex and age squared, body mass index (BMI), race, handedness, years of education, cube root of intracranial volume (ICV) and total gray matter volume (TGMV), mean frame-wise displacement. P-values were assessed via 5,000 permutation tests, and confidence intervals (CIs) were derived from 1,000 bootstrap iterations.

We further validated the heritability of INF amplitudes using cosine similarity analysis, comparing amplitude profiles between participant pairs after regressing out the same confounding variables (see Supplementary Figure 10a).

### Behavioral analysis

As in our previous study [28], we performed hierarchical factor analysis on the behavioral data in HCP S1200 using maximum likelihood estimation with orthogonal (varimax) rotation to identify latent behavioral dimensions [41]. The four-factor model was selected based on prior evidence of robustness, which includes: (1) *Mental health* – covering emotional well-being (e.g., life satisfaction, social support) as well as challenges such as anxiety and depression; (2) *Cognition* – representing intellectual abilities including memory, reasoning, and language; (3) *Processing speed* – capturing the efficiency of cognitive processing and quick responding; (4) *Substance use* – reflecting tendencies toward alcohol, tobacco, and drug use.

To assess test–retest reliability, we restricted the analysis to participants who completed both REST1 and REST2 sessions (*N* = 855). Associations between behavioral factors and average INF amplitudes were tested separately for each session using general linear models within an ANCOVA framework, controlling for age, sex, handedness, and the cube roots of ICV and TGMV. False discovery rate (FDR) was controlled using the Benjamini–Hochberg procedure.

### Spatial correlation test

To assess the statistical significance of spatial correlations between two cortical surface maps, we used the spin test, which generates null distributions by applying random rotations to the cortical sphere. Both maps were simultaneously rotated on the left and right hemisphere spherical surfaces (fsLR 32k), and Pearson correlations were recomputed for each of 1,000 permutations. P-values were calculated as the proportion of null correlations exceeding the observed correlation in absolute value.

For subcortical and cerebellar structures, where the spin test is not applicable due to the absence of a spherical surface representation, we used Moran spectral randomization. A spatial weight matrix was constructed from the inverse Euclidean distance between voxel coordinates in MNI space, and Moran eigenvectors were computed separately for left and right subdivisions of each structure (or as a single set for midline structures such as the brainstem). Surrogate maps preserving the spatial autocorrelation of the original data were generated using 1,000 permutations, and p-values were computed identically to the spin test.

Both methods were implemented using the BrainSpace toolbox [89].

## Data availability

The resting-state fMRI data analyzed in this study were obtained from the 2017 release of human connectome project (HCP) S1200 (https://www.humanconnectome.org/study/hcp-young-adult/document/1200-subjects-data-release). Since 2025, HCP has transitioned to the 2025 release as the primary distribution, and the legacy 2017 release is accessible only through Amazon S3. The multi-domain task battery (MDTB) dataset is publicly available on OpenNeuro (https://openneuro.org/datasets/ds002105). The audiovisual speech perception dataset is also publicly available on OpenNeuro (https://openneuro.org/datasets/ds003717).

## Code availability

All analyses in this study were conducted using MATLAB R2025a. The scripts and code supporting the study’s findings are available on GitHub and will be publicly accessible upon the paper’s publication.

## Author contributions

Y.S. and A.I. conceived the study. Y.S. developed the methodology, performed formal analysis and validation, created the visualizations, and wrote the original draft. J.C. and V.D.C. contributed to project development through feedback and discussion, provided resources, and reviewed and edited the manuscript. A.I. supervised the study, guided project development through feedback and direction, provided resources, and reviewed and edited the manuscript.

## Acknowledgement

This work was supported by the National Institutes of Health grant numbers R01MH136665 (to A.I. and J.C.) and R01MH123610 (to V.D.C.), and National Science Foundation grant numbers 2112455 (to V.D.C.) and 2316421 (to V.D.C.).

## References

1. Kirschner, M. and J. Gerhart, Evolvability. Proceedings of the National Academy of Sciences, 1998. 95(15): p. 8420–8427.

2. Park, H.-J. and K. Friston, Structural and functional brain networks: from connections to cognition. Science, 2013. 342(6158): p. 1238411.

3. Cole, M.W., et al., Intrinsic and task-evoked network architectures of the human brain. Neuron, 2014. 83(1): p. 238–251.

4. Gratton, C., et al., Functional brain networks are dominated by stable group and individual factors, not cognitive or daily variation. Neuron, 2018. 98(2): p. 439–452. e5.

5. Calhoun, V.D., K.A. Kiehl, and G.D. Pearlson, Modulation of temporally coherent brain networks estimated using ICA at rest and during cognitive tasks. Human brain mapping, 2008. 29(7): p. 828–838.

6. Fox, M.D., et al., The human brain is intrinsically organized into dynamic, anticorrelated functional networks. Proceedings of the National Academy of Sciences, 2005. 102(27): p. 9673–9678.

7. Spreng, R.N., et al., Default network activity, coupled with the frontoparietal control network, supports goal-directed cognition. Neuroimage, 2010. 53(1): p. 303–317.

8. Beaty, R.E., et al., Default and executive network coupling supports creative idea production. Scientific reports, 2015. 5(1): p. 10964.

9. Friston, K.J., et al., Psychophysiological and modulatory interactions in neuroimaging. Neuroimage, 1997. 6(3): p. 218–229.

10. Simony, E., et al., Dynamic reconfiguration of the default mode network during narrative comprehension. Nature communications, 2016. 7(1): p. 12141.

11. Di, X. and B.B. Biswal, Toward task connectomics: examining whole-brain task modulated connectivity in different task domains. Cerebral Cortex, 2019. 29(4): p. 1572–1583.

12. Calhoun, V.D., et al., The chronnectome: time-varying connectivity networks as the next frontier in fMRI data discovery. Neuron, 2014. 84(2): p. 262–274.

13. Lurie, D.J., et al., Questions and controversies in the study of time-varying functional connectivity in resting fMRI. Network neuroscience, 2020. 4(1): p. 30–69.

14. Iraji, A., et al., Tools of the trade: estimating time-varying connectivity patterns from fMRI data. Social cognitive and affective neuroscience, 2021. 16(8): p. 849–874.

15. Greene, A.S., et al., Why is everyone talking about brain state? Trends in Neurosciences, 2023. 46(7): p. 508–524.

16. Gu, S., et al., Controllability of structural brain networks. Nature communications, 2015. 6(1): p. 8414.

17. Parkes, L., et al., A network control theory pipeline for studying the dynamics of the structural connectome. Nature protocols, 2024. 19(12): p. 3721–3749.

18. He, B.J., Spontaneous and task-evoked brain activity negatively interact. Journal of Neuroscience, 2013. 33(11): p. 4672–4682.

19. Gao, S., G. Mishne, and D. Scheinost, Nonlinear manifold learning in functional magnetic resonance imaging uncovers a low-dimensional space of brain dynamics. Human brain mapping, 2021. 42(14): p. 4510–4524.

20. Smith, S.M., et al., Correspondence of the brain’s functional architecture during activation and rest. Proceedings of the national academy of sciences, 2009. 106(31): p. 13040–13045.

21. Abbas, A., et al., Quasi-periodic patterns contribute to functional connectivity in the brain. Neuroimage, 2019. 191: p. 193–204.

22. Yousefi, B., et al., Quasi-periodic patterns of intrinsic brain activity in individuals and their relationship to global signal. Neuroimage, 2018. 167: p. 297–308.

23. Raut, R.V., et al., Global waves synchronize the brain’s functional systems with fluctuating arousal. Science advances, 2021. 7(30): p. eabf2709.

24. Gu, Y., et al., Brain activity fluctuations propagate as waves traversing the cortical hierarchy. Cerebral cortex, 2021. 31(9): p. 3986–4005.

25. Vézquez-Rodríguez, B., et al., Signal propagation via cortical hierarchies. Network neuroscience, 2020. 4(4): p. 1072–1090.

26. Pines, A., et al., Development of top-down cortical propagations in youth. Neuron, 2023. 111(8): p. 1316–1330. e5.

27. Bolt, T., et al., A parsimonious description of global functional brain organization in three spatiotemporal patterns. Nature Neuroscience, 2022. 25(8): p. 1093–1103.

28. Song, Y., et al., Large-scale Signal Propagation Modes in the Human Brain. Neuroimage, 2025. 318: p. 121357.

29. Iraji, A., et al., Space: a missing piece of the dynamic puzzle. Trends in cognitive sciences, 2020. 24(2): p. 135–149.

30. Shine, J.M., et al., Human cognition involves the dynamic integration of neural activity and neuromodulatory systems. Nature neuroscience, 2019. 22(2): p. 289–296.

31. Song, H., W.M. Shim, and M.D. Rosenberg, Large-scale neural dynamics in a shared low-dimensional state space reflect cognitive and attentional dynamics. Elife, 2023. 12: p. e85487.

32. Margulies, D.S., et al., Situating the default-mode network along a principal gradient of macroscale cortical organization. Proceedings of the National Academy of Sciences, 2016. 113(44): p. 12574–12579.

33. Wu, X., et al., Dynamic changes in brain lateralization correlate with human cognitive performance. PLoS biology, 2022. 20(3): p. e3001560.

34. Katsumi, Y., et al., Correspondence of functional connectivity gradients across human isocortex, cerebellum, and hippocampus. Communications Biology, 2023. 6(1): p. 401.

35. Agcaoglu, O., et al., Lateralization of resting state networks and relationship to age and gender. Neuroimage, 2015. 104: p. 310–325.

36. Sridharan, D., D.J. Levitin, and V. Menon, A critical role for the right fronto-insular cortex in switching between central-executive and default-mode networks. Proceedings of the National Academy of Sciences, 2008. 105(34): p. 12569–12574.

37. Menon, V., Large-scale brain networks and psychopathology: a unifying triple network model. Trends in cognitive sciences, 2011. 15(10): p. 483–506.

38. King, M., et al., Functional boundaries in the human cerebellum revealed by a multi-domain task battery. Nature neuroscience, 2019. 22(8): p. 1371–1378.

39. Peelle, J.E., et al., Increased connectivity among sensory and motor regions during visual and audiovisual speech perception. Journal of neuroscience, 2022. 42(3): p. 435–442.

40. Chen, X., E. Viding, and T. Nichols, Faster Accelerated Permutation Inference for the ACE Model (APACE) with Parallelization. 2017.

41. Schöttner, M., et al., Exploring the latent structure of behavior using the Human Connectome Project’s data. Scientific Reports, 2023. 13(1): p. 713.

42. Tononi, G., O. Sporns, and G.M. Edelman, A measure for brain complexity: relating functional segregation and integration in the nervous system. Proceedings of the National Academy of Sciences, 1994. 91(11): p. 5033–5037.

43. Friston, K.J., Modalities, modes, and models in functional neuroimaging. Science, 2009. 326(5951): p. 399–403.

44. Sporns, O., Network attributes for segregation and integration in the human brain. Current opinion in neurobiology, 2013. 23(2): p. 162–171.

45. Chen, J.E., et al., Introducing co-activation pattern metrics to quantify spontaneous brain network dynamics. Neuroimage, 2015. 111: p. 476–488.

46. Harel, N., et al., Origin of negative blood oxygenation level—dependent fMRI signals. Journal of cerebral blood flow & metabolism, 2002. 22(8): p. 908–917.

47. Shmuel, A., et al., Negative functional MRI response correlates with decreases in neuronal activity in monkey visual area V1. Nature neuroscience, 2006. 9(4): p. 569–577.

48. Devor, A., et al., Suppressed neuronal activity and concurrent arteriolar vasoconstriction may explain negative blood oxygenation level-dependent signal. Journal of Neuroscience, 2007. 27(16): p. 4452–4459.

49. Mullinger, K.J., et al., Evidence that the negative BOLD response is neuronal in origin: a simultaneous EEG–BOLD–CBF study in humans. Neuroimage, 2014. 94: p. 263–274.

50. Logothetis, N.K., What we can do and what we cannot do with fMRI. Nature, 2008. 453(7197): p. 869–878.

51. McKiernan, K.A., et al., A parametric manipulation of factors affecting task-induced deactivation in functional neuroimaging. Journal of cognitive neuroscience, 2003. 15(3): p. 394–408.

52. Pallesen, K.J., et al., Cognitive and emotional modulation of brain default operation. Journal of Cognitive Neuroscience, 2009. 21(6): p. 1065–1080.

53. Singh, K.D. and I.P. Fawcett, Transient and linearly graded deactivation of the human default-mode network by a visual detection task. Neuroimage, 2008. 41(1): p. 100–112.

54. Gonzalez-Castillo, J., et al., Whole-brain, time-locked activation with simple tasks revealed using massive averaging and model-free analysis. Proceedings of the National Academy of Sciences, 2012. 109(14): p. 5487–5492.

55. Xu, J., V.D. Calhoun, and M.N. Potenza, The absence of task-related increases in BOLD signal does not equate to absence of task-related brain activation. Journal of neuroscience methods, 2015. 240: p. 125–127.

56. Xu, J., et al., Task-related concurrent but opposite modulations of overlapping functional networks as revealed by spatial ICA. Neuroimage, 2013. 79: p. 62–71.

57. Xu, Y., et al., Interacting spiral wave patterns underlie complex brain dynamics and are related to cognitive processing. Nature human behaviour, 2023. 7(7): p. 1196–1215.

58. Siegel, M., M.R. Warden, and E.K. Miller, Phase-dependent neuronal coding of objects in short-term memory. Proceedings of the National Academy of Sciences, 2009. 106(50): p. 21341–21346.

59. Han, H.-B., et al., Working memory readout varies with frontal theta rhythms. Neuron, 2026. 114(1): p. 159–166.e2.

60. Guell, X., et al., Functional gradients of the cerebellum. elife, 2018. 7: p. e36652.

61. Yang, S., et al., The thalamic functional gradient and its relationship to structural basis and cognitive relevance. NeuroImage, 2020. 218: p. 116960.

62. Chaudhuri, R., et al., A large-scale circuit mechanism for hierarchical dynamical processing in the primate cortex. Neuron, 2015. 88(2): p. 419–431.

63. Pang, J.C., et al., Geometric constraints on human brain function. Nature, 2023. 618(7965): p. 566–574.

64. Atasoy, S., I. Donnelly, and J. Pearson, Human brain networks function in connectome-specific harmonic waves. Nature communications, 2016. 7(1): p. 10340.

65. Jirsa, V.K. and H. Haken, Field theory of electromagnetic brain activity. Physical review letters, 1996. 77(5): p. 960.

66. Robinson, P.A., C.J. Rennie, and J.J. Wright, Propagation and stability of waves of electrical activity in the cerebral cortex. Physical Review E, 1997. 56(1): p. 826.

67. Koopman, B.O., Hamiltonian systems and transformation in Hilbert space. Proceedings of the National Academy of Sciences, 1931. 17(5): p. 315–318.

68. Kutz, J.N., et al., Dynamic mode decomposition: data-driven modeling of complex systems. 2016: SIAM.

69. Tu, J.H., et al., On dynamic mode decomposition: Theory and applications. Journal of Computational Dynamics, 2014. 1(2): p. 391–421.

70. Arbabi, H. and I. Mezic, Ergodic theory, dynamic mode decomposition, and computation of spectral properties of the Koopman operator. SIAM Journal on Applied Dynamical Systems, 2017. 16(4): p. 2096–2126.

71. O’Connor, D., et al., Identifying dynamic reproducible brain states using a predictive modelling approach. Imaging Neuroscience, 2025. 3: p. pimag_a_00540.

72. Lee, K., et al., Human brain state dynamics reflect individual neuro-phenotypes. bioRxiv, 2024: p. 2023.09. 18.557763.

73. Van Essen, D.C., et al., The WU-Minn human connectome project: an overview. Neuroimage, 2013. 80: p. 62–79.

74. Smith, S.M., et al., Resting-state fMRI in the human connectome project. Neuroimage, 2013. 80: p. 144–168.

75. Crum, R.M., et al., Population-based norms for the Mini-Mental State Examination by age and educational level. Jama, 1993. 269(18): p. 2386–2391.

76. Finn, E.S., et al., Functional connectome fingerprinting: identifying individuals using patterns of brain connectivity. Nature neuroscience, 2015. 18(11): p. 1664–1671.

77. Dubois, J., et al., A distributed brain network predicts general intelligence from resting-state human neuroimaging data. Philosophical Transactions of the Royal Society B: Biological Sciences, 2018. 373(1756): p. 20170284.

78. Marcus, D.S., et al., Human Connectome Project informatics: quality control, database services, and data visualization. Neuroimage, 2013. 80: p. 202–219.

79. Esteban, O., et al., fMRIPrep: a robust preprocessing pipeline for functional MRI. Nature methods, 2019. 16(1): p. 111–116.

80. Calhoun, V.D., et al., A method for making group inferences from functional MRI data using independent component analysis. Human brain mapping, 2001. 14(3): p. 140–151.

81. Smith, S.M., et al., Group-PCA for very large fMRI datasets. Neuroimage, 2014. 101: p. 738–749.

82. Beckmann, C.F., et al., Group comparison of resting-state FMRI data using multi-subject ICA and dual regression. Neuroimage, 2009. 47(Suppl 1): p. S148.

83. Calhoun, V.D., J.J. Pekar, and G.D. Pearlson, Alcohol intoxication effects on simulated driving: exploring alcohol-dose effects on brain activation using functional MRI. Neuropsychopharmacology, 2004. 29(11): p. 2097–2107.

84. Erhardt, E.B., et al., Comparison of multi-subject ICA methods for analysis of fMRI data. Human brain mapping, 2011. 32(12): p. 2075–2095.

85. Griffanti, L., et al., ICA-based artefact removal and accelerated fMRI acquisition for improved resting state network imaging. Neuroimage, 2014. 95: p. 232–247.

86. Pruim, R.H., et al., ICA-AROMA: A robust ICA-based strategy for removing motion artifacts from fMRI data. Neuroimage, 2015. 112: p. 267–277.

87. Dawson, S.T., et al., Characterizing and correcting for the effect of sensor noise in the dynamic mode decomposition. Experiments in Fluids, 2016. 57: p. 1–19.

88. Casorso, J., et al., Dynamic mode decomposition of resting-state and task fMRI. NeuroImage, 2019. 194: p. 42–54.

89. Vos de Wael, R., et al., BrainSpace: a toolbox for the analysis of macroscale gradients in neuroimaging and connectomics datasets. Communications biology, 2020. 3(1): p. 103.

90. Penny, W.D., et al., Statistical parametric mapping: the analysis of functional brain images. 2011: Elsevier.

91. Glasser, M.F., et al., A multi-modal parcellation of human cerebral cortex. Nature, 2016. 536(7615): p. 171–178.

92. Jovanović, M.R., P.J. Schmid, and J.W. Nichols, Sparsity-promoting dynamic mode decomposition. Physics of Fluids, 2014. 26(2).

