## Supplementary Material for "Cognition emerges from phase dynamics of intrinsic coordination"

### Supplementary Results

*Supplementary Result 1 – Predictive validity replication results*

To assess the reproducibility of the findings reported in the main text, we repeated the full predictive validation procedure in an independent replication sample drawn from the HCP dataset (n = 210; 105 training, 105 test; see Methods for sample selection). The replication sample had no overlap with the discovery sample.

All findings were reproduced (Supplementary Figure 4a). Across all tested dimensionalities (Q = 10–100), prediction performance ($R^2$) of the group-level coordination structure significantly outperformed the AR(1) null model ($p < 1.0 \times 10^{-73}$), confirming that the extracted flows capture structured dynamics beyond temporal autocorrelation. Group-level modes again yielded higher predictive accuracy than subject-specific modes across most dimensionalities ($p < 0.001$ except Q = 20). At Q = 20, the common structure still yielded higher accuracy than the subject-specific one, although this difference did not reach statistical significance.

We also evaluated prediction performance at coarse intervals (Q = 10 to 100 in steps of 10; Supplementary Figure 4b). Performance peaked at Q = 30, which significantly outperformed all other dimensionalities ($p < 0.001$). We then conducted a finer search within the range Q = 20 to 30 in steps of 1 (Supplementary Figure 4c). The highest prediction accuracy was achieved at Q = 27, corresponding to approximately 13 INF modes. Performance at Q = 28, 29, and 30 did not differ significantly from Q = 27, but Q = 27 was adopted as the optimal dimensionality for all subsequent analyses given its numerical peak.

*Supplementary Result 2 – Reproducibility of INF modes*

To assess the reproducibility of the estimated INF modes, we independently applied the full INF mode estimation procedure (with $Q = 27$ INs) to the REST1 (n = 951) and REST2 (n = 884) sessions of the HCP dataset. Mode correspondence between sessions was evaluated using the modal assurance criterion (MAC; Pastor et al., 2012), which quantifies the similarity between two complex-valued mode vectors as the squared magnitude of their normalized inner product (Supplementary Figure 5). Values above 0.9 indicate consistent correspondence and values below 0.6 suggest poor resemblance (Vacher et al., 2010).

INF modes 1–8 were consistently reproduced across sessions, with all diagonal MAC values exceeding 0.80; modes 1–4 and 7 showed particularly high reproducibility (MAC > 0.90). Modes 9–10 and 11–12, however, exhibited elevated off-diagonal MAC values, indicating partial mixing between mode pairs. To determine whether this reflects rotation within a shared subspace rather than a genuine failure of reproducibility, we computed the S2MAC (D'Ambrogio & Fregolent, 2003), which evaluates whether a single mode from one session can be adequately represented within the subspace spanned by a pair of modes from the other session. The S2MAC values confirmed subspace-level consistency: REST2 modes 9 and 10 achieved S2MAC values of 0.74 and 0.91 with the REST1 mode 9–10 subspace, and REST2 modes 11 and 12 achieved 0.85 and 0.87 with the REST1 mode 11–12 subspace. These results suggest that modes within each pair share similar eigenvalues, leading to rotational ambiguity in the eigen-decomposition, while the subspaces they span—and thus the dynamics they generate—remain consistent across sessions. Mode 13 showed lower reproducibility (MAC = 0.36) and warrants further investigation in future work.

Overall, the core INF modes (1–8) demonstrated high test–retest reproducibility, and modes 9–12 were consistent at the subspace level, supporting the robustness of the estimated coordination structure.

*Supplementary Result 3 – Derivation of closed-form FC from INF modes*

Under the INF framework, assuming no amplitude decay, the whole-brain signal at time $t$, given an initial INF phase $\theta_k$ at $t = 0$, becomes:

$$\boldsymbol{x}(t) = \sum_k 2\,\mathrm{Re}\big[\eta_k e^{i\theta_k} e^{i\omega_k t} S\boldsymbol{\psi}_k\big]$$

where $\eta_k$ is INF amplitude, $\omega_k$ is the intrinsic evolution rate of mode $k$, $S$ denotes IN spatial maps, and $\boldsymbol{\psi}_k$ is INF mode k. Then, the signal at region $i$ is modeled as a superposition of cosine components (since $\mathrm{Re}\big[e^{i\theta}\big] = \cos\theta$):

$$x_i(t) = \sum_k \alpha_{ik} \cos\left(\omega_k t + \theta_k + \gamma_{ik}\right)$$

where $\alpha_{ik}$ is the regional magnitude at region $i$ in mode $k$ (which includes the effects of conjugate vector and INF amplitude), $\gamma_{ik} = \arg[(S\boldsymbol{\psi}_k)_i]$ is the phase lag (flow sequence position) of region $i$ in mode $k$. Note that the INF phase $\theta_k$ acts as a global offset common to all regions within mode $k$.

The Pearson correlation between regions $i$ and $j$ is defined as:

$$\rho_{ij} = \frac{\langle x_i(t)\, x_j(t)\rangle}{\sqrt{\langle x_i(t)^2\rangle \langle x_j(t)^2\rangle}}$$

where $\langle\cdot\rangle$ denotes the long-time average, $\langle f(t)\rangle = \lim_{T\to\infty} \frac{1}{2T}\int_{-T}^{T} f(t)\, dt$.

**Numerator.** Expanding the product:

$$\big\langle x_i(t)\, x_j(t)\big\rangle = \sum_k \sum_l \alpha_{ik}\alpha_{jl} \big\langle \cos(\omega_k t + \theta_k + \gamma_{ik}) \cos\big(\omega_l t + \theta_l + \gamma_{jl}\big)\big\rangle$$

Applying the product-to-sum identity:

$$\cos A \cos B = \frac{1}{2}\left[\cos\left(A - B\right) + \cos\left(A + B\right)\right]$$

each term becomes:

$$\frac{\alpha_{ik}\,\alpha_{jl}}{2}\big[\langle\cos\left((\omega_k-\omega_l)t+(\theta_k-\theta_l)+(\gamma_{ik}-\gamma_{jl})\right)\rangle+\langle\cos\left((\omega_k+\omega_l)t+(\theta_k+\theta_l)\right.$$
$$\left.+(\gamma_{ik}+\gamma_{jl})\right)\rangle\big]$$

Under long-time averaging, any term containing a non-zero frequency argument vanishes:

$$\langle\cos(\omega t+c)\rangle=0\ \text{ for }\ \omega\neq 0$$

The second term always vanishes since $\omega_k+\omega_l>0$. The first term survives only when $\omega_k=\omega_l$, i.e., $k=l$ (assuming distinct evolution rates across modes). In this case:

$$\langle\cos\left(\theta_k-\theta_l+\gamma_{ik}-\gamma_{jk}\right)\rangle=\cos\left(\gamma_{ik}-\gamma_{jk}\right)$$

since $k=l$. Therefore:

$$\langle x_i(t)\,x_j(t)\rangle=\frac{1}{2}\sum_k\alpha_{ik}\alpha_{jk}\cos\left(\gamma_{ik}-\gamma_{jk}\right)$$

**Denominator.** Setting $i=j$ in the above:

$$\langle x_i(t)^2\rangle=\frac{1}{2}\sum_k\alpha_{ik}^2\cos\left(0\right)=\frac{1}{2}\sum_k\alpha_{ik}^2$$

and likewise for region $j$.

**Result.** Combining numerator and denominator:

$$\rho_{ij}=\frac{\frac{1}{2}\sum_k\alpha_{ik}\alpha_{jk}\cos\left(\gamma_{ik}-\gamma_{jk}\right)}{\sqrt{\frac{1}{2}\sum_k\alpha_{ik}^2\cdot\frac{1}{2}\sum_k\alpha_{jk}^2}}=\frac{\sum_k\alpha_{ik}\,\alpha_{jk}\cos\left(\gamma_{ik}-\gamma_{jk}\right)}{\sqrt{\sum_k\alpha_{ik}^2\cdot\sum_k\alpha_{jk}^2}}$$

Critically, the INF phase $\theta_k$ does not appear in the final expression: because $\theta_k$ is a global offset shared by all regions within mode $k$, it cancels in the pairwise spatial phase difference. The reconstructed FC is therefore invariant under any phase modulation $\theta_k\rightarrow\theta_k+\Delta\theta_k$. This provides the theoretical guarantee that FC topology is preserved across cognitive states within the INF framework.

**Supplementary Figures**

*Supplementary Figure 1*

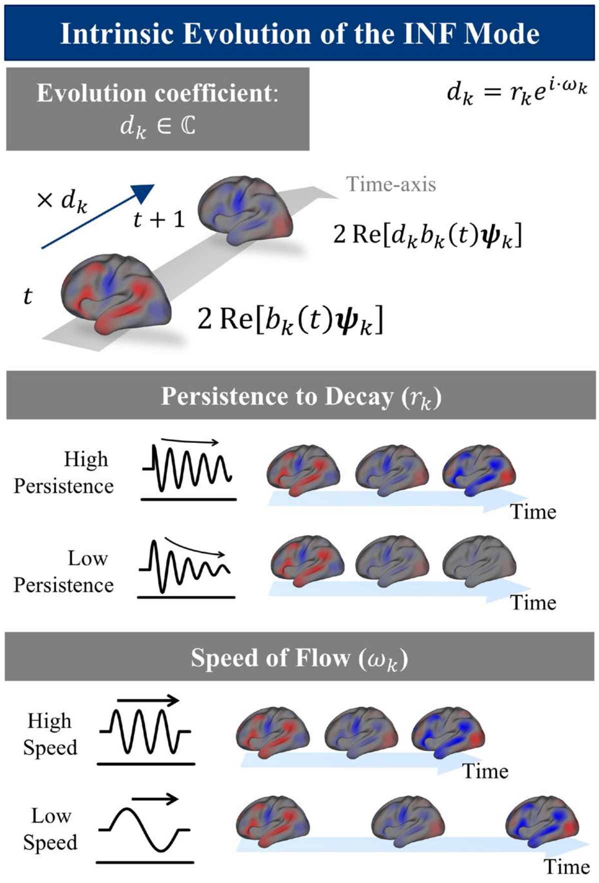


**Supplementary Figure 1. Intrinsic temporal evolution of INF modes.** In the absence of internal or external task demands, each flow coefficient evolves autonomously through multiplication by a complex-valued evolution coefficient $d_k = r_k e^{i\omega_k}$, such that $b_k(t+1) = d_k \cdot b_k(t)$. This single scalar captures two properties of the flow's intrinsic dynamics. The persistence rate ($r_k$) governs how rapidly the flow amplitude decays over time: high persistence ($r_k \approx 1$) maintains a stable oscillation, whereas low persistence leads to rapid attenuation. The speed of flow ($\omega_k$) determines how quickly the INF phase advances per time step, and thus how rapidly the signal propagates through the flow sequence: high speed produces fast temporal progression across networks, whereas low speed yields slow progression.

*Supplementary Figure 2*

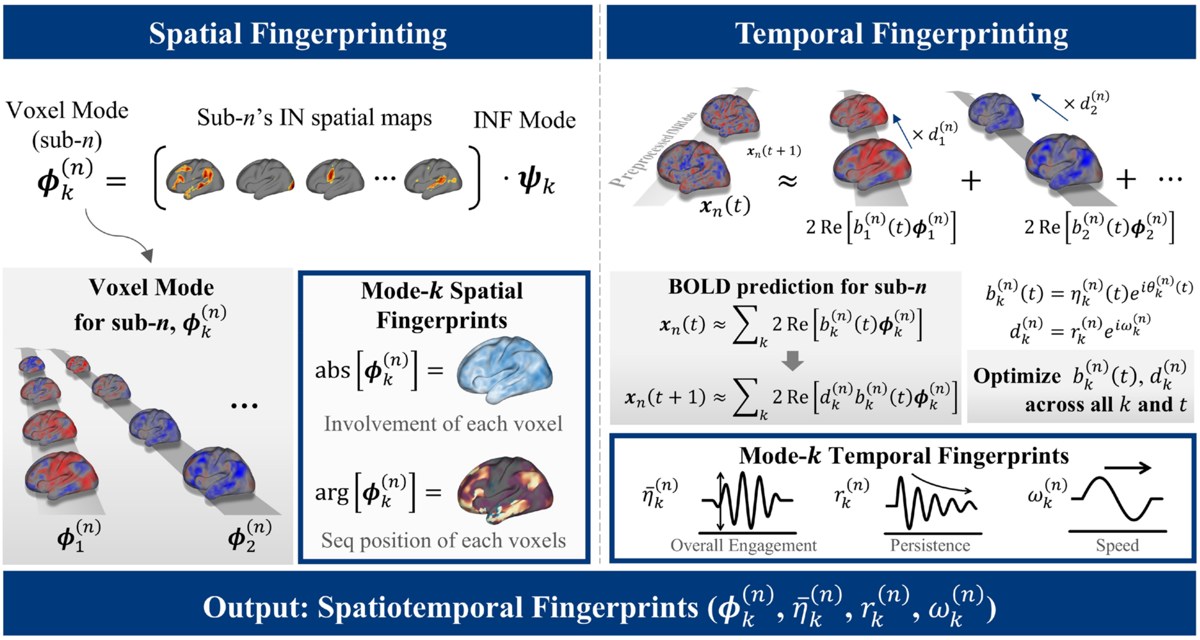


**Supplementary Figure 2. Spatiotemporal fingerprinting.** *Spatial fingerprinting*: each INF mode $\boldsymbol{\psi}_k$ is projected into individual subjects' BOLD space via subject-specific IN spatial maps, producing voxel-level mode expressions $\boldsymbol{\phi}_k^{(n)}$, referred to as spatial fingerprints. The magnitude of each voxel reflects its involvement in the mode; the argument reflects its sequential position within the flow. *Temporal fingerprinting*: subject-specific temporal parameters—mean flow amplitude ($\bar{\eta}_k^{(n)}$) and intrinsic evolution rate ($d_k^{(n)}$, which further decomposed into persistence rate $r_k^{(n)}$, and flow speed $\omega_k^{(n)}$)—are estimated by optimizing these parameters to best predict future BOLD signals (see Methods). Together, each subject's spatial and temporal fingerprints constitute their unique spatiotemporal fingerprint. See Methods for implementation details.

*Supplementary Figure 3*

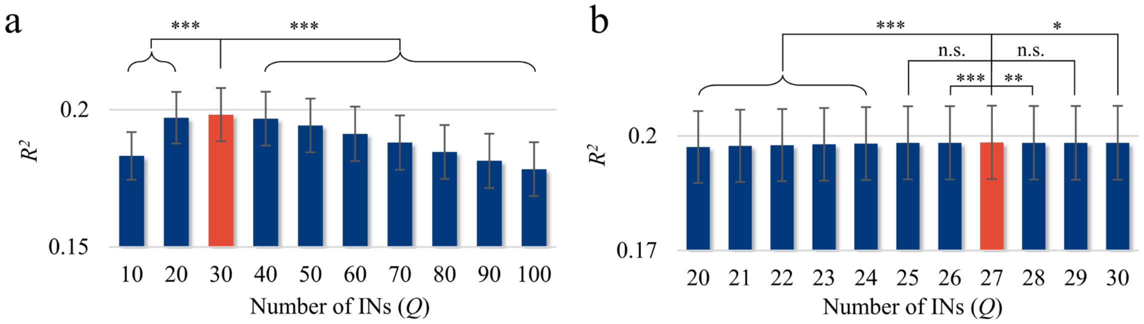


**Supplementary Figure 3. Dimensionality selection and fingerprinting contribution. (a)** Prediction performance ($R^2$) within the group-level coordination structure across a coarse range of IN dimensionalities (Q = 10–100 in steps of 10). **(b)** Finer-grained search within Q = 20–30 (steps of 1). The best-performing dimensionality (Q = 27, highlighted in orange) was adopted for all subsequent analyses, although performance did not differ significantly from that at Q = 25 or Q = 29. This similarity among odd dimensionalities (25, 27, and 29) may reflect the importance of the non-flow mode (i.e., modes that capture stationary spatial patterns rather than flow-like dynamics), as an odd dimensionality guarantees at least one real eigenvector (see Methods).

*Supplementary Figure 4.*

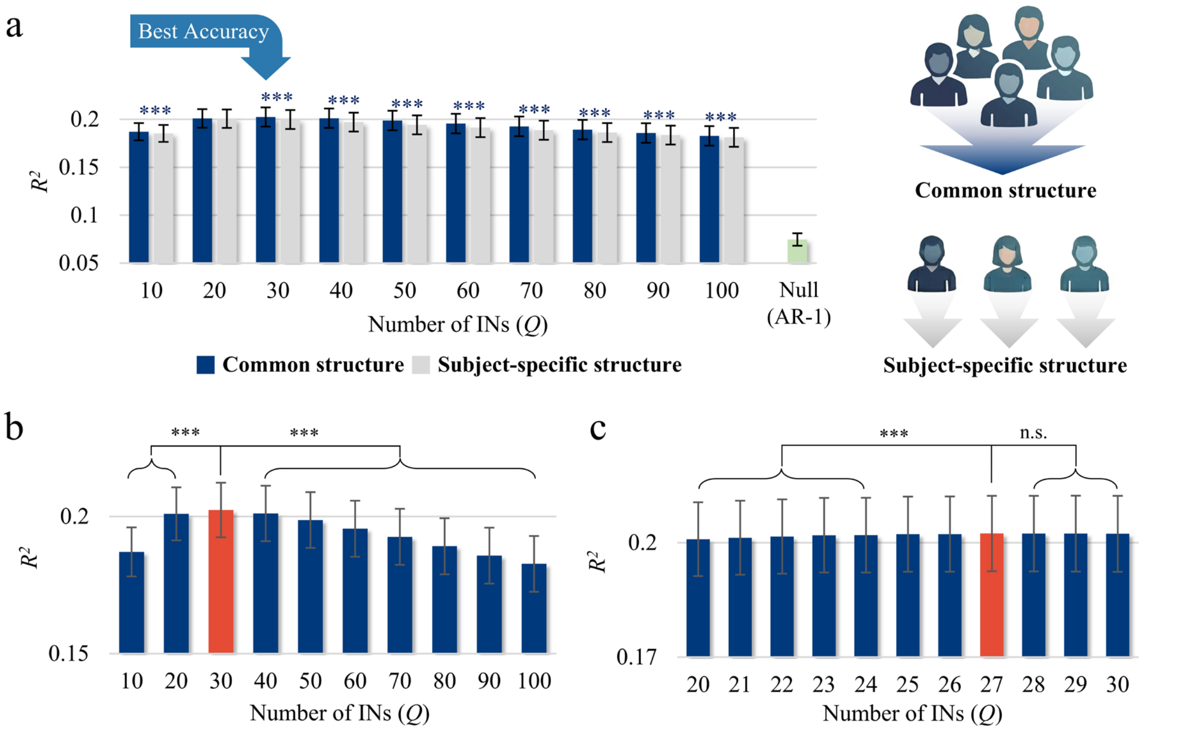


**Supplementary Figure 4. Replication of predictive validation in the replication set (n = 210). (a)** Prediction performance ($R^2$) using a group-level (common) coordination structure (i.e., group-level INF modes; dark blue) versus subject-specific structures (i.e., subject-specific INF modes; grey) across IN dimensionalities (Q = 10–100), with the AR(1) null model shown for reference (green). Results replicate those of the discovery sample (Figure 3): the common structure significantly outperformed both the null model and subject-specific structures across most dimensionalities. At Q = 20, the common structure still yielded higher accuracy than the subject-specific one, although this difference did not reach statistical significance. **(b)** Prediction performance within the group-level coordination structure across a coarse range of dimensionalities (Q = 10–100 in steps of 10). Performance peaked at Q = 30. **(c)** Finer-grained search within Q = 20–30 (steps of 1). The highest accuracy was achieved at Q = 27 (highlighted in orange), consistent with the discovery sample, although performance did not differ significantly from Q = 25 and 28–30.

*Supplementary Figure 5*

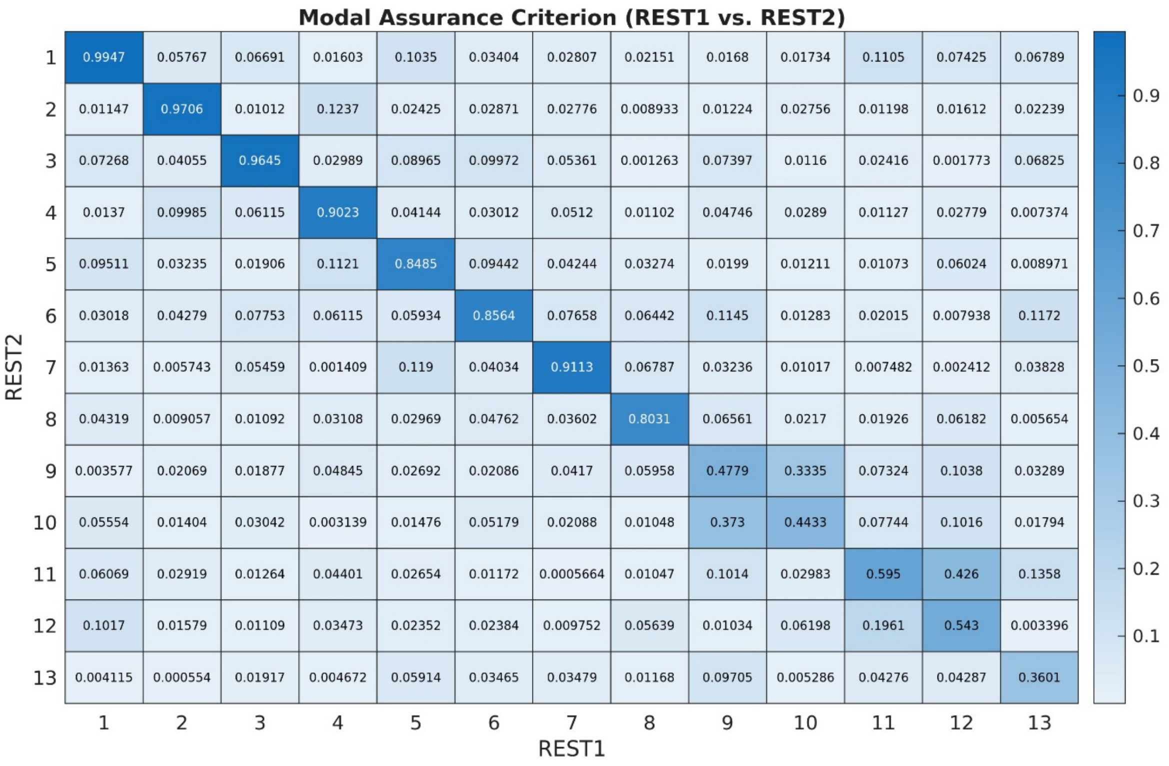


**Supplementary Figure 5. INF mode reproducibility results.** Modal assurance criterion (MAC) matrix comparing INF modes estimated independently from the HCP REST1 (n = 951) and REST2 (n = 884) sessions. Each element represents the squared magnitude of the normalized inner product between a REST1 mode (columns) and a REST2 mode (rows). High diagonal values indicate consistent mode recovery across sessions.

*Supplementary Figure 6*

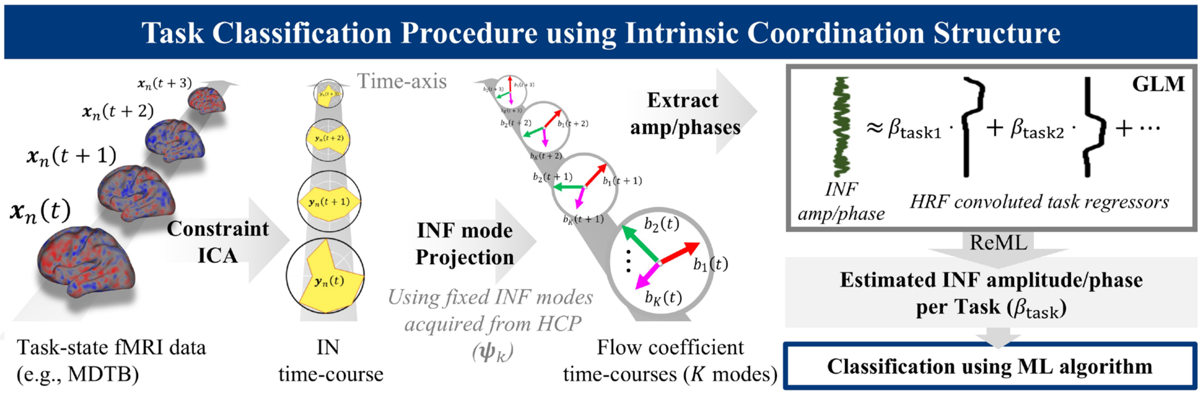


**Supplementary Figure 6. Task classification procedure using INF.** Task-state fMRI data are first projected onto group-level IN spatial maps via constrained ICA using template spatial maps (estimated from HCP resting-state data), ensuring that the resulting IN spatial maps correspond to those used to define the INF mode vectors. These time-courses are then projected onto the fixed INF mode vectors ($\boldsymbol{\psi}_k$, estimated from HCP resting-state data) to yield flow coefficient time courses $b_k(t)$. Then, INF amplitude and phase are extracted from these complex-valued coefficients; the phase is further decomposed into cosine and sine components to enable linear modeling. Separate general linear models (GLMs) with HRF-convolved task regressors and AR(1) temporal autocorrelation are then performed via restricted maximum likelihood (ReML) for each feature type—amplitude, $\cos(\theta)$, and $\sin(\theta)$. This yields condition-specific beta values ($\beta_{\text{task}}$) for each feature. These beta values serve as input features for task classification using a machine learning algorithm. See Methods for implementation details.

*Supplementary Figure 7*

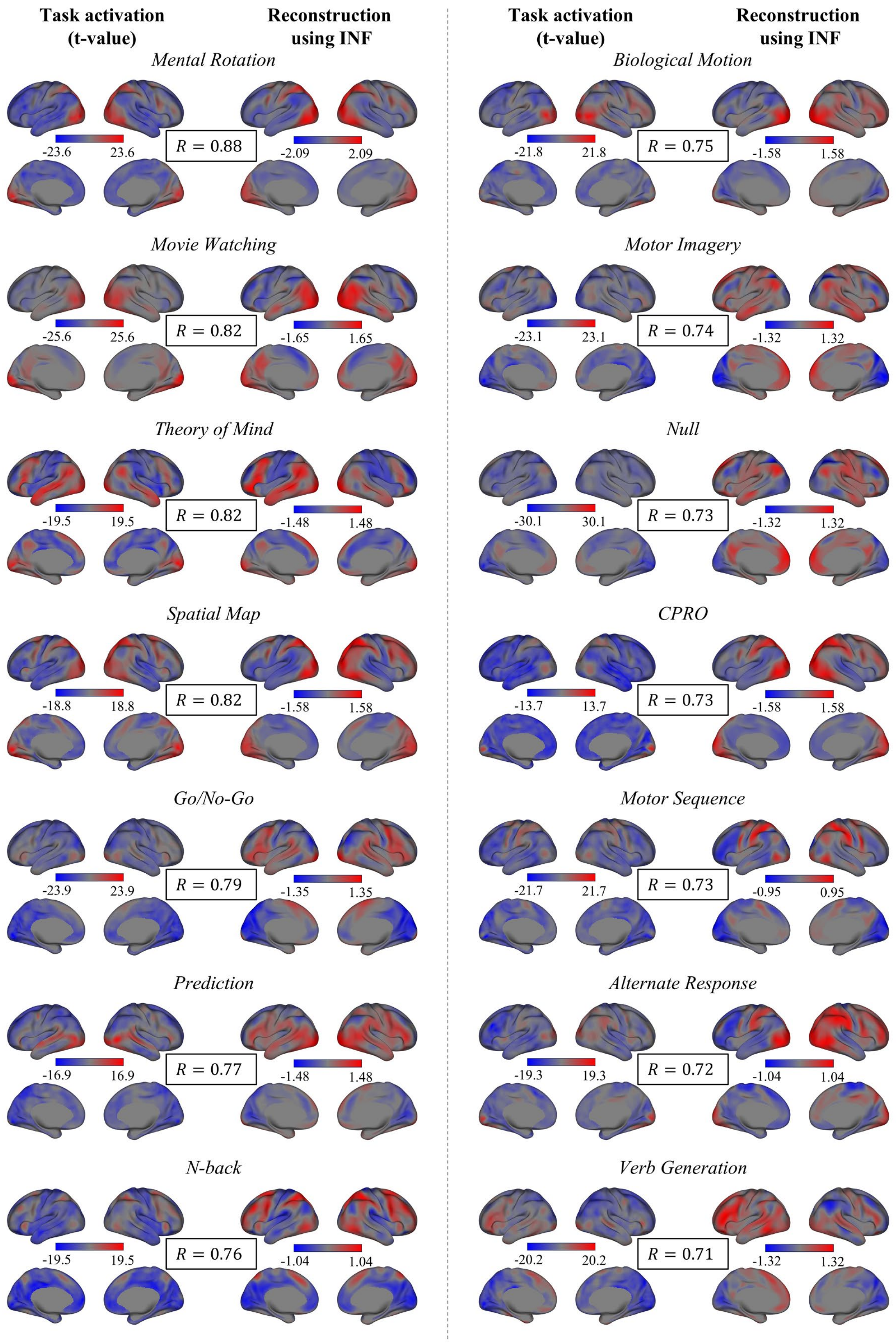

*Supplementary Figure 7 (cont.)*

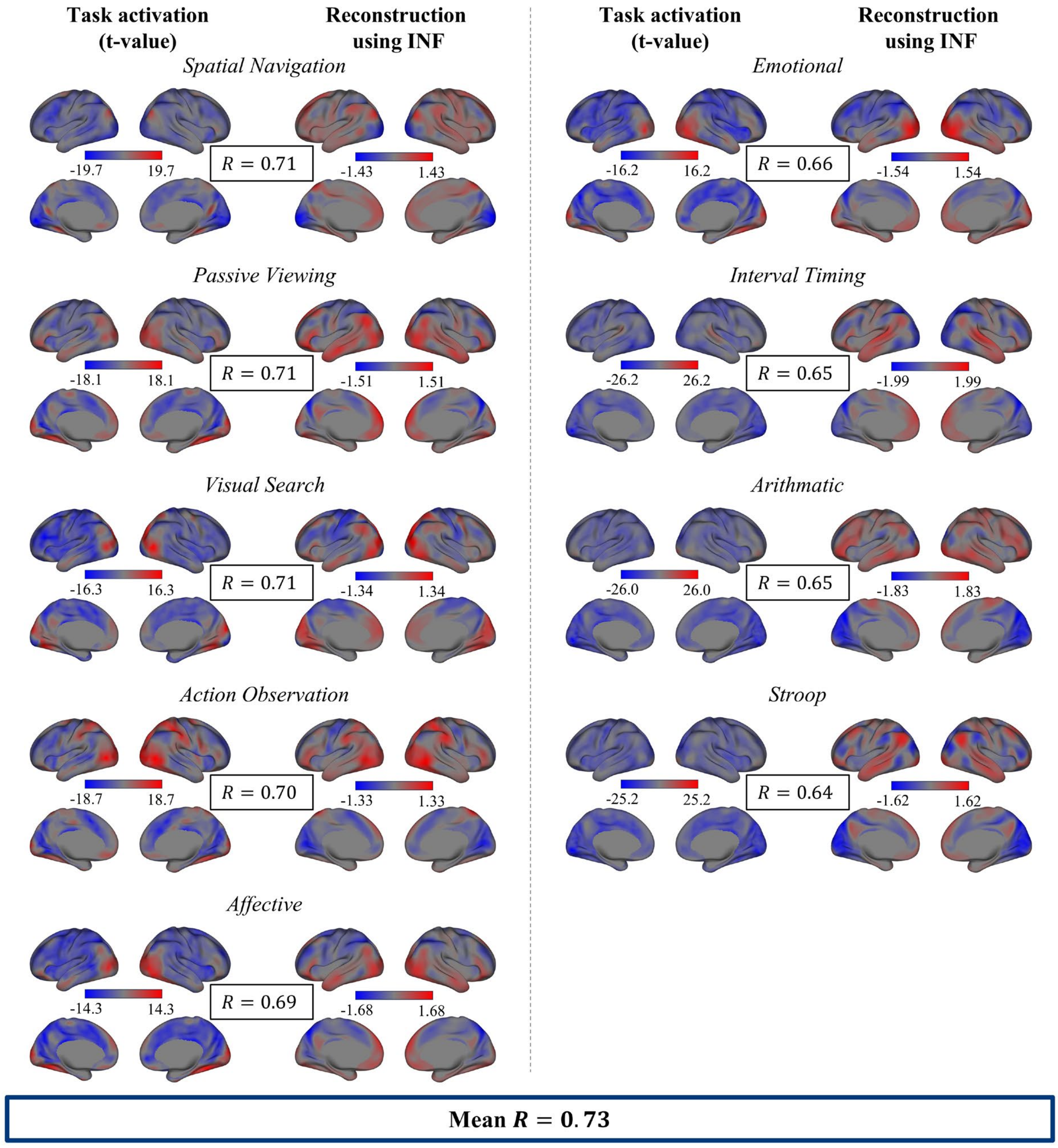


**Supplementary Figure 7. Task-evoked activation reconstruction for all 23 MDTB tasks.** Empirical task activation maps (t-value; left) and their reconstructions using INF phase modulation (right) for all 23 tasks, sorted by spatial correlation in descending order.

*Supplementary Figure 8*

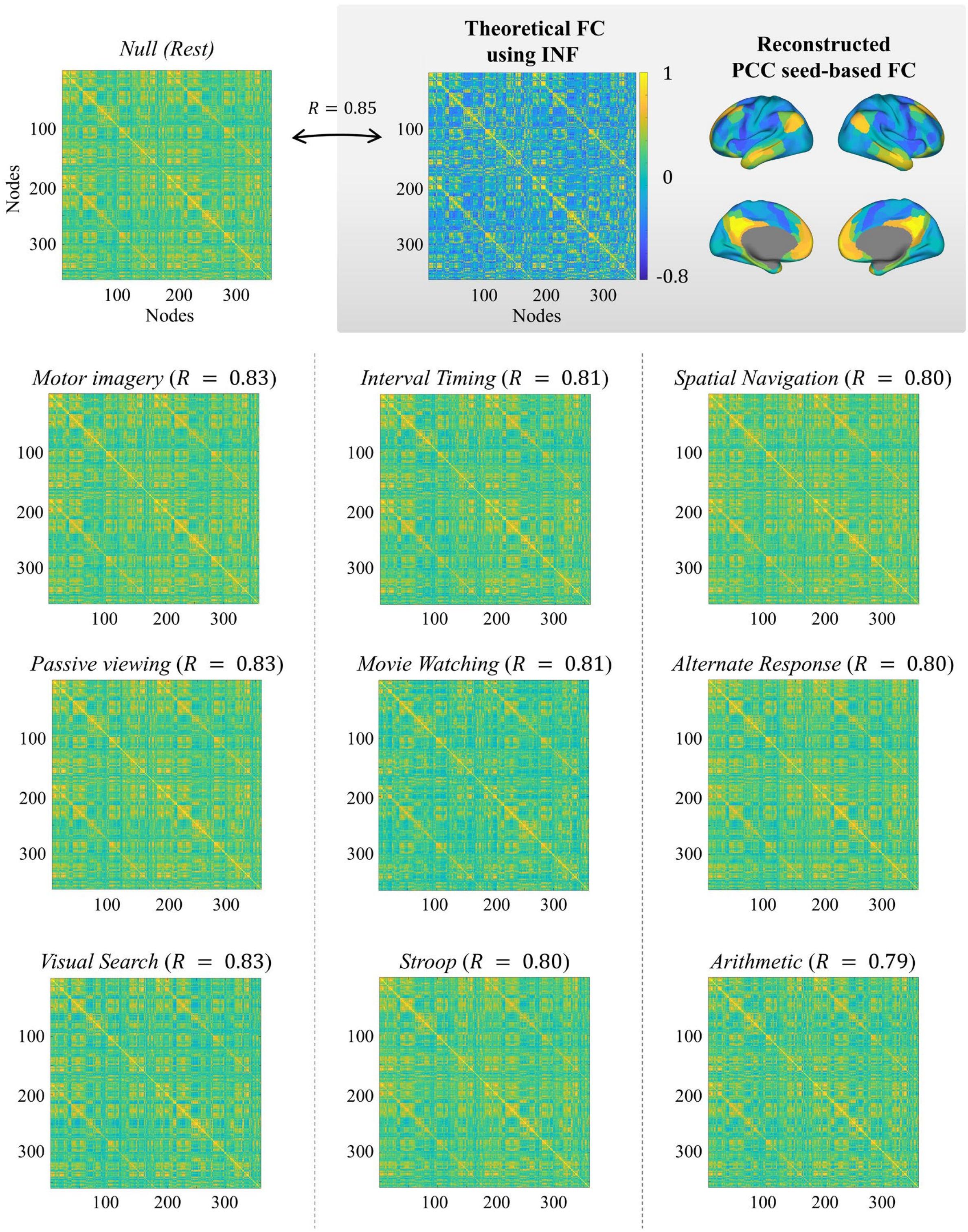

*Supplementary Figure 8 (cont.)*

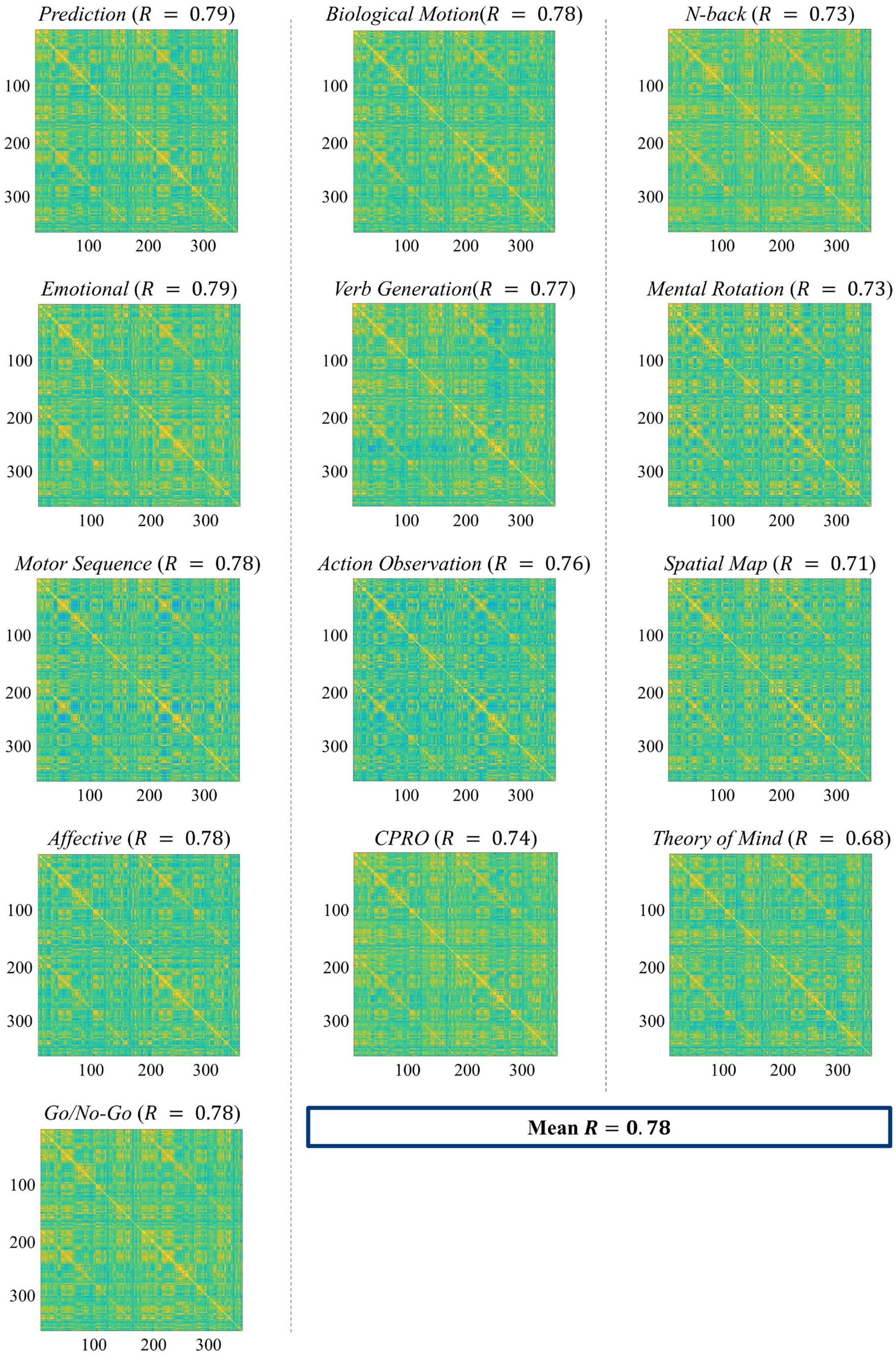

**Supplementary Figure 8. FC matrices for all 23 MDTB tasks and comparison with INF-reconstructed FC.** FC matrices were computed from each task condition and compared with the theoretical FC reconstructed from resting-state INF parameters. The grey inset shows the INF-reconstructed FC matrix (left) and reconstructed PCC seed-based FC map (right), the latter illustrating the anti-correlation structure between DMN and CEN. Top left: the FC matrix from the rest condition, which shows strong correspondence with the INF-reconstructed FC (R = 0.85). Below: FC matrices from all remaining task conditions, displayed in descending order of similarity (Pearson's R) to the INF-reconstructed FC. The correlation value is indicated next to each task name.

*Supplementary Figure 9*

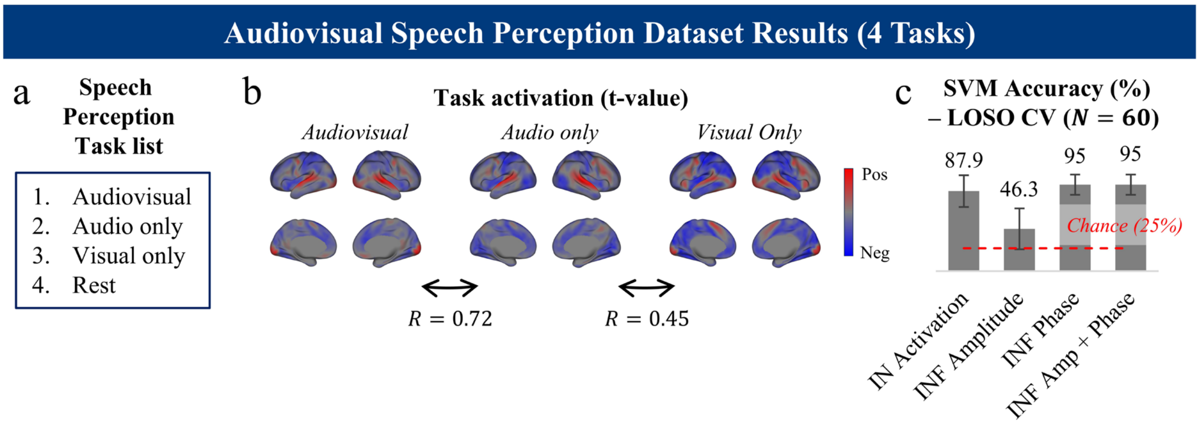


**Supplementary Figure 9. Task classification in the audiovisual speech perception dataset. (a)** List of four task conditions. **(b)** Group-level task-evoked activation maps (t-value) for the three speech perception conditions. Activation maps were highly similar across conditions (in particular, audiovisual vs. audio only: R = 0.72), posing a challenging classification problem using activation patterns. **(c)** Leave-one-subject-out (LOSO) classification accuracy (N = 60) using a linear SVM across four feature types. Red dashed line indicates chance level (25%). INF phase substantially outperformed activation-based features, demonstrating that phase captures task-state distinctions even when activation maps are highly overlapping. Error bars denote standard deviations.

*Supplementary Figure 10*

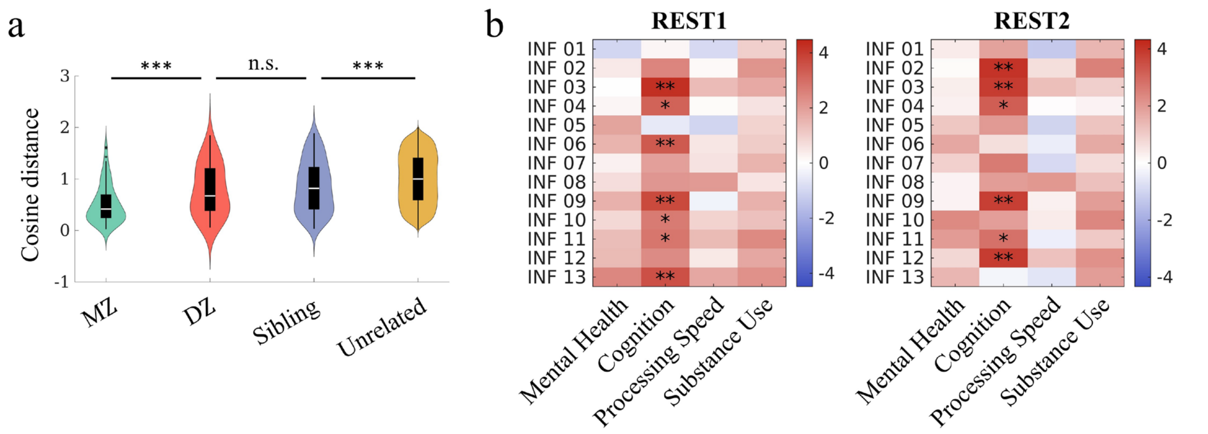


**Supplementary Figure 10. Heritability and behavioral associations of INF amplitudes. (a)** Cosine similarity of mean INF amplitude profiles between pairs of participants, computed after regressing out confounding variables (age, age squared, sex, the interaction between age and sex, the interaction between sex and age squared, BMI, race, handedness, years of education, cube root of ICV and TGMV, mean frame-wise displacement). Similarity was compared across four kinship categories: monozygotic twins (MZ), dizygotic twins (DZ), non-twin siblings, and unrelated pairs using two-sample t-tests (MZ vs. DZ: $p = 2.79 \times 10^{-5}$; DZ vs. sibling: $p = 0.096$; sibling vs. unrelated: $p = 5.57 \times 10^{-13}$). The graded decrease in similarity with decreasing genetic relatedness supports a heritable component of INF amplitude profiles. **(b)** T-value heatmaps for correlation between mean INF amplitudes and behavioral scores (see Methods). INF mode estimation, fingerprinting, and behavioral correlation were performed independently for REST1 and REST2 sessions to assess reproducibility. Across both sessions, INF mode amplitudes showed overall positive associations with the cognition factor, which reflects general intellectual ability. INF modes 3, 4, 9, and 11 exhibit consistent correlations across sessions, although modes 9 and 11 shows only moderate similarity between REST1 and REST2. Asterisks indicate significance after FDR correction (*q<0.05, **q<0.01).

**Supplementary Videos 1–13.**

Temporal evolution of INF modes 1–13, showing reconstructed activity patterns across cortical, subcortical, and cerebellar regions. Each frame advances the INF phase by $\Delta\theta = \pi/30$, displaying one full oscillation cycle ($[0, 2\pi]$, 60 frames) per mode.

**Supplementary Methods**

*Supplementary Method 1 – fMRIPrep preprocessing details for MDTB dataset*

Results included in this manuscript come from preprocessing performed using *fMRIPrep* 25.2.3 (Esteban et al. (2019); Esteban et al. (2018); RRID:SCR_016216), which is based on *Nipype* 1.10.0 (K. Gorgolewski et al. (2011); K. J. Gorgolewski et al. (2018); RRID:SCR_002502).

Preprocessing of $B_0$ inhomogeneity mappings

A total of 4 fieldmaps were found available within the input BIDS structure for this particular subject. A $B_0$ nonuniformity map (or *fieldmap*) was estimated from the phase-drift map(s) measure with two consecutive GRE (gradient-recalled echo) acquisitions. The corresponding phase-map(s) were phase-unwrapped with prelude (FSL None).

Anatomical data preprocessing

A total of 1 T1-weighted (T1w) images were found within the input BIDS dataset. The T1w image was corrected for intensity non-uniformity (INU) with N4BiasFieldCorrection (Tustison et al. 2010), distributed with ANTs 2.6.2 (Avants et al. 2008, RRID:SCR_004757), and used as T1w-reference throughout the workflow. The T1w-reference was then skull-stripped with a *Nipype* implementation of the antsBrainExtraction.sh workflow (from ANTs), using OASIS30ANTs as target template. Brain tissue segmentation of cerebrospinal fluid (CSF), white-matter (WM) and gray-matter (GM) was performed on the brain-extracted T1w using fast (FSL (version unknown), RRID:SCR_002823, Zhang, Brady, and Smith 2001). Brain surfaces were reconstructed using recon-all (FreeSurfer 7.3.2, RRID:SCR_001847, Dale, Fischl, and Sereno 1999), and the brain mask estimated previously was refined with a custom variation of the method to reconcile ANTs-derived and FreeSurfer-derived segmentations of the cortical gray-matter of Mindboggle (RRID:SCR_002438, Klein et al. 2017). A T2-weighted image was used to improve pial surface refinement. Volume-based spatial normalization to two standard spaces (MNI152NLin2009cAsym, MNI152NLin6Asym) was performed through nonlinear registration with antsRegistration (ANTs 2.6.2), using brain-extracted versions of both T1w reference and the T1w template. The following templates were were selected for spatial normalization and accessed with *TemplateFlow* (25.0.4, Ciric et al. 2022): *ICBM 152 Nonlinear Asymmetrical template version 2009c* [Fonov et al. (2009), RRID:SCR_008796; TemplateFlow ID: MNI152NLin2009cAsym], *FSL's MNI ICBM 152 non-linear 6th Generation Asymmetric Average Brain Stereotaxic Registration Model* [Evans et al. (2012), RRID:SCR_002823; TemplateFlow ID: MNI152NLin6Asym].

Functional data preprocessing

For each of the 34 BOLD runs found per subject (across all tasks and sessions), the following preprocessing was performed. First, a reference volume was generated, using a custom methodology of *fMRIPrep*, for use in head motion correction. Head-motion parameters with respect to the BOLD reference (transformation matrices, and six corresponding rotation and translation parameters) are estimated before any spatiotemporal filtering using mcflirt (FSL , Jenkinson et al. 2002). The estimated *fieldmap* was then aligned with rigid-registration to the target EPI (echo-planar imaging) reference run. The field coefficients were mapped on to the reference EPI using the transform. The BOLD reference was then co-registered to the T1w reference using bbregister (FreeSurfer) which implements boundary-based registration (Greve and Fischl 2009). Co-registration was configured with six degrees of freedom. The aligned T2w image was used for initial co-registration.Several confounding time-series were calculated based on the *preprocessed BOLD*: framewise displacement (FD), DVARS and three region-wise global signals. FD was computed using two formulations following Power (absolute sum of relative motions, Power et al. (2014)) and Jenkinson (relative root mean square displacement between affines, Jenkinson et al. (2002)). FD and DVARS are calculated for each functional run, both using their implementations in *Nipype* (following the definitions by Power et al. 2014). The three global signals are extracted within the CSF, the WM, and the whole-brain masks. Additionally, a set of physiological regressors were extracted to allow for component-based noise correction (*CompCor*, Behzadi et al. 2007). Principal components are estimated after high-pass filtering the *preprocessed BOLD* time-series (using a discrete cosine filter with 128s cut-off) for the two *CompCor* variants: temporal (tCompCor) and anatomical (aCompCor). tCompCor components are then calculated from the top 2% variable voxels within the brain mask. For aCompCor, three probabilistic masks (CSF, WM and combined CSF+WM) are generated in anatomical space. The implementation differs from that of Behzadi et al. in that instead of eroding the masks by 2 pixels on BOLD space, a mask of pixels that likely contain a volume fraction of GM is subtracted from the aCompCor masks. This mask is obtained by dilating a GM mask extracted from the FreeSurfer's *aseg* segmentation, and it ensures components are not extracted from voxels containing a minimal fraction of GM. Finally, these masks are resampled into BOLD space and binarized by thresholding at 0.99 (as in the original implementation). Components are also calculated separately within the WM and CSF masks. For each CompCor decomposition, the *k* components with the largest singular values are retained, such that the retained components' time series are sufficient to explain 50 percent of variance across the nuisance mask (CSF, WM, combined, or temporal). The remaining components are dropped from consideration. The head-motion estimates calculated in the correction step were also placed within the corresponding confounds file. The confound time series derived from head motion estimates and global signals were expanded with the inclusion of temporal derivatives and quadratic terms for each (Satterthwaite et al. 2013). Frames that exceeded a threshold of 0.5 mm FD or 1.5 standardized DVARS were annotated as motion outliers. Additional nuisance timeseries are calculated by means of principal components analysis of the signal found within a thin band (*crown*) of voxels around the edge of the brain, as proposed by (Patriat, Reynolds, and Birn 2017). The BOLD time-series were resampled onto the left/right-symmetric template “fsLR” using the Connectome Workbench (Glasser et al. 2013). *Grayordinates* files (Glasser et al. 2013) containing 91k samples were also generated with surface data transformed directly to fsLR space and subcortical data transformed to 2 mm resolution MNI152NLin6Asym space. All resamplings can be performed with *a single interpolation step* by composing all the pertinent transformations (i.e. head-motion transform matrices, susceptibility distortion correction when available, and co-registrations to anatomical and output spaces). Gridded (volumetric) resamplings were performed using nitransforms, configured with cubic B-spline interpolation.

Functional data preprocessing

For each of the 34 BOLD runs found per subject (across all tasks and sessions), the following preprocessing was performed. First, a reference volume was generated, using a custom methodology of *fMRIPrep*, for use in head motion correction. Head-motion parameters with respect to the BOLD reference (transformation matrices, and six corresponding rotation and translation parameters) are estimated before any spatiotemporal filtering using mcflirt (FSL , Jenkinson et al. 2002). The BOLD reference was then co-registered to the T1w reference using bbregister (FreeSurfer) which implements boundary-based registration (Greve and Fischl 2009). Co-registration was configured with six degrees of freedom. The aligned T2w image was used for initial co-registration.Several confounding time-series were calculated based on the *preprocessed BOLD*: framewise displacement (FD), DVARS and three region-wise global signals. FD was computed using two formulations following Power (absolute sum of relative motions, Power et al. (2014)) and Jenkinson (relative root mean square displacement between affines, Jenkinson et al. (2002)). FD and DVARS are calculated for each functional run, both using their implementations in *Nipype* (following the definitions by Power et al. 2014). The three global signals are extracted within the CSF, the WM, and the whole-brain masks. Additionally, a set of physiological regressors were extracted to allow for component-based noise correction (*CompCor*, Behzadi et al. 2007). Principal components are estimated after high-pass filtering the *preprocessed BOLD* time-series (using a discrete cosine filter with 128s cut-off) for the two *CompCor* variants: temporal (tCompCor) and anatomical (aCompCor). tCompCor components are then calculated from the top 2% variable voxels within the brain mask. For aCompCor, three probabilistic masks (CSF, WM and combined CSF+WM) are generated in anatomical space. The implementation differs from that of Behzadi et al. in that instead of eroding the masks by 2 pixels on BOLD space, a mask of pixels that likely contain a volume fraction of GM is subtracted from the aCompCor masks. This mask is obtained by dilating a GM mask extracted from the FreeSurfer's *aseg* segmentation, and it ensures components are not extracted from voxels containing a minimal fraction of GM. Finally, these masks are resampled into BOLD space and binarized by thresholding at 0.99 (as in the original implementation). Components are also calculated separately within the WM and CSF masks. For each CompCor decomposition, the *k* components with the largest singular values are retained, such that the retained components' time series are sufficient to explain 50 percent of variance across the nuisance mask (CSF, WM, combined, or temporal). The remaining components are dropped from consideration. The head-motion estimates calculated in the correction step were also placed within the corresponding confounds file. The confound time series derived from head motion estimates and global signals were expanded with the inclusion of temporal derivatives and quadratic terms for each (Satterthwaite et al. 2013). Frames that exceeded a threshold of 0.5 mm FD or 1.5 standardized DVARS were annotated as motion outliers. Additional nuisance timeseries are calculated by means of principal components analysis of the signal found within a thin band (*crown*) of voxels around the edge of the brain, as proposed by (Patriat, Reynolds, and Birn 2017). The BOLD time-series were resampled onto the left/right-symmetric template "fsLR" using the Connectome Workbench (Glasser et al. 2013). *Grayordinates* files (Glasser et al. 2013) containing 91k samples were also generated with surface data transformed directly to fsLR space and subcortical data transformed to 2 mm resolution MNI152NLin6Asym space. All resamplings can be performed with *a single interpolation step* by composing all the pertinent transformations (i.e. head-motion transform matrices, susceptibility distortion correction when available, and co-registrations to anatomical and output spaces). Gridded (volumetric) resamplings were performed using nitransforms, configured with cubic B-spline interpolation. *Grayordinate* "dscalar" files containing 91k samples were resampled onto fsLR using the Connectome Workbench (Glasser et al. 2013). Many internal operations of *fMRIPrep* use *Nilearn* 0.11.1 (Abraham et al. 2014, RRID:SCR_001362), mostly within the functional processing workflow. For more details of the pipeline, see the section corresponding to workflows in *fMRIPrep*'s documentation.

Copyright Waiver

The above boilerplate text was automatically generated by fMRIPrep with the express intention that users should copy and paste this text into their manuscripts *unchanged*. It is released under the CC0 license.

*Supplementary Method 2 – fMRIPrep preprocessing details for audiovisual speech perception dataset*

Results included in this manuscript come from preprocessing performed using *fMRIPrep* 25.2.3 (Esteban et al. (2019); Esteban et al. (2018); RRID:SCR_016216), which is based on *Nipype* 1.10.0 (K. Gorgolewski et al. (2011); K. J. Gorgolewski et al. (2018); RRID:SCR_002502).

Anatomical data preprocessing

A total of 1 T1-weighted (T1w) images were found within the input BIDS dataset. The T1w image was corrected for intensity non-uniformity (INU) with N4BiasFieldCorrection (Tustison et al. 2010), distributed with ANTs 2.6.2 (Avants et al. 2008, RRID:SCR_004757), and used as T1w-reference throughout the workflow. The T1w-reference was then skull-stripped with a *Nipype* implementation of the antsBrainExtraction.sh workflow (from ANTs), using OASIS30ANTs as target template. Brain tissue segmentation of cerebrospinal fluid (CSF), white-matter (WM) and gray-matter (GM) was performed on the brain-extracted T1w using fast (FSL (version unknown), RRID:SCR_002823, Zhang, Brady, and Smith 2001). Brain surfaces were reconstructed using recon-all (FreeSurfer 7.3.2, RRID:SCR_001847, Dale, Fischl, and Sereno 1999), and the brain mask estimated previously was refined with a custom variation of the method to reconcile ANTs-derived and FreeSurfer-derived segmentations of the cortical gray-matter of Mindboggle (RRID:SCR_002438, Klein et al. 2017). A T2-weighted image was used to improve pial surface refinement. Brain surfaces were reconstructed using recon-all (FreeSurfer 7.3.2, RRID:SCR_001847, Dale, Fischl, and Sereno 1999), and the brain mask estimated previously was refined with a custom variation of the method to reconcile ANTs-derived and FreeSurfer-derived segmentations of the cortical gray-matter of Mindboggle (RRID:SCR_002438, Klein et al. 2017). Volume-based spatial normalization to two standard spaces (MNI152NLin2009cAsym, MNI152NLin6Asym) was performed through nonlinear registration with antsRegistration (ANTs 2.6.2), using brain-extracted versions of both T1w reference and the T1w template. The following templates were were selected for spatial normalization and accessed with *TemplateFlow* (25.0.4, Ciric et al. 2022): *ICBM 152 Nonlinear Asymmetrical template version 2009c* [Fonov et al. (2009), RRID:SCR_008796; TemplateFlow ID: MNI152NLin2009cAsym], *FSL's MNI ICBM 152 non-linear 6th Generation Asymmetric Average Brain Stereotaxic Registration Model* [Evans et al. (2012), RRID:SCR_002823; TemplateFlow ID: MNI152NLin6Asym].

Functional data preprocessing

For each of the 6 BOLD runs found per subject (across all tasks and sessions), the following preprocessing was performed. First, a reference volume was generated, using a custom methodology of *fMRIPrep*, for use in head motion correction. Head-motion parameters with respect to the BOLD reference (transformation matrices, and six corresponding rotation and translation parameters) are estimated before any spatiotemporal filtering using mcflirt (FSL , Jenkinson et al. 2002). The BOLD reference was then co-registered to the T1w reference using bbregister (FreeSurfer) which implements boundary-based registration (Greve and Fischl 2009). Co-registration was configured with six degrees of freedom. The aligned T2w image was used for initial co-registration.Several confounding time-series were calculated based on the *preprocessed BOLD*: framewise displacement (FD), DVARS and three region-wise global signals. FD was computed using two formulations following Power (absolute sum of relative motions, Power et al. (2014)) and Jenkinson (relative root mean square displacement between affines, Jenkinson et al. (2002)). FD and DVARS are calculated for each functional run, both using their implementations in *Nipype* (following the definitions by Power et al. 2014). The three global signals are extracted within the CSF, the WM, and the whole-brain masks. Additionally, a set of physiological regressors were extracted to allow for component-based noise correction (*CompCor*, Behzadi et al. 2007). Principal components are estimated after high-pass filtering the *preprocessed BOLD* time-series (using a discrete cosine filter with 128s cut-off) for the two *CompCor* variants: temporal (tCompCor) and anatomical (aCompCor). tCompCor components are then calculated from the top 2% variable voxels within the brain mask. For aCompCor, three probabilistic masks (CSF, WM and combined CSF+WM) are generated in anatomical space. The implementation differs from that of Behzadi et al. in that instead of eroding the masks by 2 pixels on BOLD space, a mask of pixels that likely contain a volume fraction of GM is subtracted from the aCompCor masks. This mask is obtained by dilating a GM mask extracted from the FreeSurfer's *aseg* segmentation, and it ensures components are not extracted from voxels containing a minimal fraction of GM. Finally, these masks are resampled into BOLD space and binarized by thresholding at 0.99 (as in the original implementation). Components are also calculated separately within the WM and CSF masks. For each CompCor decomposition, the *k* components with the largest singular values are retained, such that the retained components' time series are sufficient to explain 50 percent of variance across the nuisance mask (CSF, WM, combined, or temporal). The remaining components are dropped from consideration. The head-motion estimates calculated in the correction step were also placed within the corresponding confounds file. The confound time series derived from head motion estimates and global signals were expanded with the inclusion of temporal derivatives and quadratic terms for each (Satterthwaite et al. 2013). Frames that exceeded a threshold of 0.5 mm FD or 1.5 standardized DVARS were annotated as motion outliers. Additional nuisance timeseries are calculated by means of principal components analysis of the signal found within a thin band (*crown*) of voxels around the edge of the brain, as proposed by (Patriat, Reynolds, and Birn 2017). The BOLD time-series were resampled onto the left/right-symmetric template "fsLR" using the Connectome Workbench (Glasser et al.

2013). *Grayordinates* files (Glasser et al. 2013) containing 91k samples were also generated with surface data transformed directly to fsLR space and subcortical data transformed to 2 mm resolution MNI152NLin6Asym space. All resamplings can be performed with *a single interpolation step* by composing all the pertinent transformations (i.e. head-motion transform matrices, susceptibility distortion correction when available, and co-registrations to anatomical and output spaces). Gridded (volumetric) resamplings were performed using nitransforms, configured with cubic B-spline interpolation. *Grayordinate* "dscalar" files containing 91k samples were resampled onto fsLR using the Connectome Workbench (Glasser et al. 2013). Many internal operations of *fMRIPrep* use *Nilearn* 0.11.1 (Abraham et al. 2014, RRID:SCR_001362), mostly within the functional processing workflow. For more details of the pipeline, see the section corresponding to workflows in *fMRIPrep*'s documentation.

Copyright Waiver

The above boilerplate text was automatically generated by fMRIPrep with the express intention that users should copy and paste this text into their manuscripts *unchanged*. It is released under the CC0 license.

**Supplementary References**


- Abraham, Alexandre, Fabian Pedregosa, Michael Eickenberg, Philippe Gervais, Andreas Mueller, Jean Kossaifi, Alexandre Gramfort, Bertrand Thirion, and Gael Varoquaux. 2014. "Machine Learning for Neuroimaging with Scikit-Learn." *Frontiers in Neuroinformatics* 8. https://doi.org/10.3389/fninf.2014.00014.
- Avants, B. B., C. L. Epstein, M. Grossman, and J. C. Gee. 2008. "Symmetric Diffeomorphic Image Registration with Cross-Correlation: Evaluating Automated Labeling of Elderly and Neurodegenerative Brain." *Medical Image Analysis* 12 (1): 26–41. https://doi.org/10.1016/j.media.2007.06.004.
- Behzadi, Yashar, Khaled Restom, Joy Liau, and Thomas T. Liu. 2007. "A Component Based Noise Correction Method (CompCor) for BOLD and Perfusion Based fMRI." *NeuroImage* 37 (1): 90–101. https://doi.org/10.1016/j.neuroimage.2007.04.042.
- Ciric, R., William H. Thompson, R. Lorenz, M. Goncalves, E. MacNicol, C. J. Markiewicz, Y. O. Halchenko, et al. 2022. "TemplateFlow: FAIR-Sharing of Multi-Scale, Multi-Species Brain Models." *Nature Methods* 19: 1568–71. https://doi.org/10.1038/s41592-022-01681-2.
- Dale, Anders M., Bruce Fischl, and Martin I. Sereno. 1999. "Cortical Surface-Based Analysis: I. Segmentation and Surface Reconstruction." *NeuroImage* 9 (2): 179–94. https://doi.org/10.1006/nimg.1998.0395.
- D'ambrogio, W., and Fregolent, A. 2003. "Higher-order mac for the correlation of close and multiple modes." Mechanical systems and signal processing, 17(3), 599-610. https://doi.org/10.1006/mssp.2002.1468
- Esteban, Oscar, Ross Blair, Christopher J. Markiewicz, Shoshana L. Berleant, Craig Moodie, Feilong Ma, Ayse Ilkay Isik, et al. 2018. "fMRIPrep 25.2.3." *Software*. https://doi.org/10.5281/zenodo.852659.

- Esteban, Oscar, Christopher Markiewicz, Ross W Blair, Craig Moodie, Ayse Ilkay Isik, Asier Erramuzpe Aliaga, James Kent, et al. 2019. “fMRIPrep: A Robust Preprocessing Pipeline for Functional MRI.” *Nature Methods* 16: 111–16. https://doi.org/10.1038/s41592-018-0235-4.
- Evans, AC, AL Janke, DL Collins, and S Baillet. 2012. “Brain Templates and Atlases.” *NeuroImage* 62 (2): 911–22. https://doi.org/10.1016/j.neuroimage.2012.01.024.
- Fonov, VS, AC Evans, RC McKinstry, CR Almli, and DL Collins. 2009. “Unbiased Nonlinear Average Age-Appropriate Brain Templates from Birth to Adulthood.” *NeuroImage* 47, Supplement 1: S102. https://doi.org/10.1016/S1053-8119(09)70884-5.
- Glasser, Matthew F., Stamatios N. Sotiropoulos, J. Anthony Wilson, Timothy S. Coalson, Bruce Fischl, Jesper L. Andersson, Junqian Xu, et al. 2013. “The Minimal Preprocessing Pipelines for the Human Connectome Project.” *NeuroImage*, Mapping the connectome, 80: 105–24. https://doi.org/10.1016/j.neuroimage.2013.04.127.
- Gorgolewski, K., C. D. Burns, C. Madison, D. Clark, Y. O. Halchenko, M. L. Waskom, and S. Ghosh. 2011. “Nipype: A Flexible, Lightweight and Extensible Neuroimaging Data Processing Framework in Python.” *Frontiers in Neuroinformatics* 5: 13. https://doi.org/10.3389/fninf.2011.00013.
- Gorgolewski, Krzysztof J., Oscar Esteban, Christopher J. Markiewicz, Erik Ziegler, David Gage Ellis, Michael Philipp Notter, Dorota Jarecka, et al. 2018. “Nipype.” *Software*. https://doi.org/10.5281/zenodo.596855.
- Greve, Douglas N, and Bruce Fischl. 2009. “Accurate and Robust Brain Image Alignment Using Boundary-Based Registration.” *NeuroImage* 48 (1): 63–72. https://doi.org/10.1016/j.neuroimage.2009.06.060.

- Jenkinson, Mark, Peter Bannister, Michael Brady, and Stephen Smith. 2002. "Improved Optimization for the Robust and Accurate Linear Registration and Motion Correction of Brain Images." *NeuroImage* 17 (2): 825–41. https://doi.org/10.1006/nimg.2002.1132.
- Klein, Arno, Satrajit S. Ghosh, Forrest S. Bao, Joachim Giard, Yrjö Häme, Eliezer Stavsky, Noah Lee, et al. 2017. "Mindboggling Morphometry of Human Brains." *PLOS Computational Biology* 13 (2): e1005350. https://doi.org/10.1371/journal.pcbi.1005350.
- Pastor, M., Binda, M., and Harčarik, T. 2012. "Modal assurance criterion." *Procedia Engineering*, *48*, 543-548. https://doi.org/10.1016/j.proeng.2012.09.551.
- Patriat, Rémi, Richard C. Reynolds, and Rasmus M. Birn. 2017. "An Improved Model of Motion-Related Signal Changes in fMRI." *NeuroImage* 144, Part A (January): 74–82. https://doi.org/10.1016/j.neuroimage.2016.08.051.
- Power, Jonathan D., Anish Mitra, Timothy O. Laumann, Abraham Z. Snyder, Bradley L. Schlaggar, and Steven E. Petersen. 2014. "Methods to Detect, Characterize, and Remove Motion Artifact in Resting State fMRI." *NeuroImage* 84 (Supplement C): 320–41. https://doi.org/10.1016/j.neuroimage.2013.08.048.
- Satterthwaite, Theodore D., Mark A. Elliott, Raphael T. Gerraty, Kosha Ruparel, James Loughead, Monica E. Calkins, Simon B. Eickhoff, et al. 2013. "An improved framework for confound regression and filtering for control of motion artifact in the preprocessing of resting-state functional connectivity data." *NeuroImage* 64 (1): 240–56. https://doi.org/10.1016/j.neuroimage.2012.08.052.
- Tustison, N. J., B. B. Avants, P. A. Cook, Y. Zheng, A. Egan, P. A. Yushkevich, and J. C. Gee. 2010. "N4ITK: Improved N3 Bias Correction." *IEEE Transactions on Medical Imaging* 29 (6): 1310–20. https://doi.org/10.1109/TMI.2010.2046908.
- Vacher, P., Jacquier, B., & Bucharles, A. 2010. "Extensions of the MAC criterion to complex modes." In *Proceedings of the international conference on noise and vibration*

*engineering* (pp. 2713-2726). ISMA Leuven, Belgium.

- Zhang, Y., M. Brady, and S. Smith. 2001. “Segmentation of Brain MR Images Through a Hidden Markov Random Field Model and the Expectation-Maximization Algorithm.” *IEEE Transactions on Medical Imaging* 20 (1): 45–57. https://doi.org/10.1109/42.906424.
